# Mechanism of manganese dysregulation of dopamine neuronal activity

**DOI:** 10.1101/792143

**Authors:** Min Lin, Luis M. Colon-Perez, Danielle O. Sambo, Douglas R. Miller, Joseph J. Lebowitz, Felix Jimenez-Rondan, Robert J. Cousins, Nicole Horenstein, Tolunay Beker Aydemir, Marcelo Febo, Habibeh Khoshbouei

**Affiliations:** Department of Neuroscience, University of Florida, Gainesville, FL 32611; Department of Psychiatry, University of Florida, Gainesville, FL 32611; Center for Nutritional Sciences, University of Florida, Gainesville, FL 32611; Department of Chemistry, University of Florida, Gainesville, FL 32611; Division of Nutritional Sciences, Cornell University, Ithaca, NY 14850

**Keywords:** Manganese, Dopamine Neurons, L-type Calcium Channel, Synaptic Transmission, Parkinson’s Disease

## Abstract

Manganese exposure produces Parkinson’s-like neurological symptoms, suggesting a selective dysregulation of dopamine transmission. It is unknown, however, how manganese accumulates in dopaminergic brain regions or how it regulates the activity of dopamine neurons. Our *in vivo* studies suggest manganese accumulates in dopamine neurons of the ventral tegmental area and substantia nigra via nifedipine-sensitive Ca^2+^ channels. Manganese produces a Ca^2+^ channel-mediated current which increases neurotransmitter release and rhythmic firing activity of dopamine neurons. These increases are prevented by blockade of Ca^2+^ channels and depend on downstream recruitment of Ca^2+^-activated potassium channels to the plasma membrane. These findings demonstrate the mechanism of manganese-induced dysfunction of dopamine neurons, and reveal a potential therapeutic target to attenuate manganese-induced impairment of dopamine transmission.

**Significance Statement:** Manganese is a trace element critical to many physiological processes. Overexposure to manganese is an environmental risk factor for neurological disorders such as a Parkinson’s disease-like syndrome known as manganism. We found manganese dose-dependently increased the excitability of dopamine neurons, decreased the amplitude of action potentials, and narrowed action potential width. Blockade of Ca^2+^ channels prevented these effects as well as manganese accumulation in the mouse midbrain *in vivo*. Our data provide a potential mechanism for manganese-regulation of dopaminergic neurons.

## Introduction

Manganese is a trace element critical to many physiological and developmental processes, including the regulation of macronutrient metabolism, blood glucose, cellular energy, reproduction, digestion, and bone growth (Greene and Madgwick, 1988; Erikson et al., 2005). Manganese is a cofactor for several enzymatic processes and a constituent of metalloenzymes, including arginase, pyruvate carboxylase, and manganese-containing superoxide dismutase (Ashner and Aschner, 2005; Ashner et al., 2007; Guilarte, 2010). Except in children on long-term parenteral nutrition or individuals with mutations in the metal transporter SLC39A8 gene, manganese deficiencies are seldom reported (Greene and Madgwick, 1988; Zogzas and Mukhopadhyay, 2017). In contrast, excess manganese accumulation in the brain following environmental exposure is implicated in abnormalities related to the dopaminergic system, including Parkinson-like motor dysfunction (Jankovic, 2005), ataxia (Soriano et al., 2016), and hallucinations (Verhoeven et al., 2011). Animal models of manganism have shown that a single large exposure or prolonged moderate exposure to excess manganese is detrimental to the basal ganglia function (Michalke and Fernsebner 2014; Olanow, 2004), albeit with less understood mechanisms.

Manganese can enter the central nervous system (CNS) through the cerebral spinal fluid or by crossing cerebral capillary endothelial membranes (Aschner et al., 2007). Physiological concentrations of manganese in the human brain range from 20 to 53 µM (Bowman and Aschner, 2014) but can increase several-fold upon overexposure both in humans (Crossgrove and Zheng, 2004; Kessler et al., 2003) as well as rodents (Liu et al., 2006). Existing studies have used a wide range (60–150 µM) of extracellular manganese to investigate manganese-associated neurotoxicity (Bowman and Aschner, 2014; Tuschl et al., 2013). Studies on manganese transport in mammalian systems have largely focused on influx mechanisms (Au et al., 2008). Manganese is transported into neurons, and possibly other CNS cell types, through a number of transporters, including divalent metal transporters (Gunshin et al., 1997), the transferrin receptor (Gunter et al., 2013), store-operated Ca^2+^ channels (Crossgrove and Yokel, 2005), the choline transporter (Lockman et al., 2001), the magnesium transporter (Goytain et al., 2008), and the NMDA receptor (Itoh et al., 2008). In addition, ZIP14 is shown to uptake manganese in human neuroblastoma cells (Fujishiro et al., 2014). The clinical effects of manganese toxicity are primarily Parkinson-like in nature (Jankovic 2005). This includes movement disorders characterized by tremor, rigidity, dystonia and/or ataxia; psychiatric disturbances including irritability, impulsiveness, agitation, obsessive-compulsive behavior, and hallucinations; and cognitive deficits such as memory impairment, reduced learning capacity, decreased mental flexibility, and cognitive slowing (Josephs et al., 2005). Neuronal degeneration and altered neurotransmitter release occur in brain regions with abnormally high accumulation of manganese, including the dorsal striatum, internal globus pallidus (GPi), and substantia nigra pars reticulata (SNpr) (Crossgrove and Zheng, 2004; Guilarte, 2010; Perl and Olanow, 2007; Uchino et al., 2007). In addition, neuronal loss and gliosis in the globus pallidus, SNpr, and striatum are reported with high accumulation of manganese (Olanow, 2004). Consistently, there is severe cell loss in the substantia nigra pars compacta (SNpc) of individuals with documented chronic manganese exposure, typically through occupational or environmental means (Perl and Olanow, 2007). More recently, mutations in the human metal transporter genes ZNT10 and ZIP14 have shown to cause manganese overload and motor dysfunction (Leyva-Illades et al., 2014; Tuschl et al., 2013). In addition to these clinical findings, previous studies show a correlation between elevated extracellular manganese levels in the brain and dysfunction of dopamine transmission (Dodd et al., 2013; Madison et al., 2012), where manganese reduced dopamine uptake and amphetamine-induced dopamine efflux (Roth et al., 2013). Manganese exposure in developing rats reduces both the levels and activity of striatal D_2_ receptors (Seth and Chandra, 1984; Rogers et al., 2014), supporting the overarching hypothesis for manganese-mediated dysregulation of the dopaminergic system.

The mechanism by which manganese dysregulates dopamine neurons is poorly understood. One potential mechanism by which manganese can regulate cellular responses is through the modulation of Ca^2+^ concentrations, which not only regulates membrane potential but also serves as an important signaling molecule (Clapham, 2007). Manganese has been shown to regulate Ca^2+^ signaling in primary astrocytes cultures where exposure to manganese results in mitochondrial sequestration of Ca^2+^ which in turn reduces the available pool of releasable Ca^2+^ within the endoplasmic reticulum (Tjalkens et al., 2006). This can affect the production of reactive oxygen species, free radicals, and toxic metabolites; alteration of mitochondrial function and ATP production; and depletion of cellular antioxidant defense mechanisms (Martinez-Finley et al., 2013; Puskin and Gunter, 1973). In primary astrocytes, manganese rapidly inhibits ATP-induced Ca^2+^ waves and Ca^2+^ transients (Streifel et al., 2013) as well as decreases the influx of extracellular Ca^2+^ induced by 1-oleoyl-2-acetyl-sn-glycerol (OAG), a direct activator of the transient receptor potential channel TRPC3 (Streifel et al., 2013).

The mechanistic relationship between manganese-regulation of Ca^2+^ signaling and excitability of dopamine neurons has remained unclear. Recently, we have shown changes in Ca^2+^ homeostasis in the dopamine neurons influence neuronal activity indirectly through Ca^2+^-activated potassium channels (Lin et al., 2016). In the current study, we report a mechanistic link between manganese regulation of the excitability of dopamine neurons and manganese modulation of Ca^2+^ channels. Here, we identified a cellular mechanism by which manganese accumulates in the midbrain, influxes into dopamine neurons, and regulates the activity of dopamine neurons. Contrary to our initial hypothesis, we found manganese does not block Ca^2+^ channels in dopamine neurons but acts as a substrate for Ca^2+^ channels. The influx of manganese through the nifedipine-sensitive Ca^2+^ channels was further supported by computational modeling, single neuron analysis, and *in vivo* magnetic resonance imaging experiments showing blockade of Cav1.2 channels decreased manganese regulation of dopaminergic neuronal activity and its accumulation in the midbrain. These data address the existing debate in the field regarding manganese regulation of Ca^2+^ channels in dopamine neurons. The mechanistic results reported here provide a clinically relevant therapeutic target that could attenuate the severity of manganese toxicity in patients exposed to excess manganese.

## Materials and Methods

### Drugs and reagents

The drugs and reagents used in this study were purchased from Millipore Sigma-Aldrich (St. Louis, Mo), unless otherwise stated. Chemical reagents and drugs used for primary neuronal culture are listed in Table 1. Antibodies used for Western blot analysis are listed in Table 2. The catalog number of the reagents and drugs used for electrophysiology and microscopy experiments are listed in Table 3.

### Animals

Midbrain neuronal cultures were obtained from RFP::TH C57BL/6 mice (obtained from Dr. Douglas McMahon, Vanderbilt University), a transgenic mouse strain where the dopamine neurons are rendered fluorescent by expressing the red fluorescent protein (RFP) under the tyrosine hydroxylase (TH) promoter (Zhang et al., 2004). Mice conditionally expressing GCaMP6f in dopaminergic neurons were generated by crossing animals expressing Cre recombinase under control of the Slc6a3 promoter (B6.SJL-Slc6a3tm1.1 (cre)BkmnI J; Jackson Lab, Stock: 006660) to animals expressing GcaMP6f under control of the LoxP promoter (B6;129s-Gt(ROSA) 26Sortm95.1 (CAG-GCaMP6f HzeI J; Jackson Lab, stock: 024105). Mice were housed in the animal care facilities at the University of Florida in accordance with Institutional Animal Care and Use Committee, under guidelines established by National Institutes of Health. Food and water were available *ad libitum* in the home cage. Animals were housed under standard conditions at 22−24°C, 50–60% humidity, and a 12 h light/dark cycle.

### Manganese concentrations used in this study

The physiological concentration of manganese in the human brain is estimated to be between 5.32 and 14.03 ng manganese /mg protein, equivalent to 20.0–52.8 μM (Bowman and Aschner, 2014). Average cellular manganese content in animal models of chronic manganese exposure is as high as 10.95 μg g^−1^ (200 μM), which was shown to decrease the viability of dopamine neurons (Higashi et al., 2004). Using this information, we first performed dose response experiments (Figure 1 and 2) and found that 100 μM manganese is a suitable concentration to study manganese-regulation of intrinsic firing behavior of dopamine neurons.

**Figure 1.**
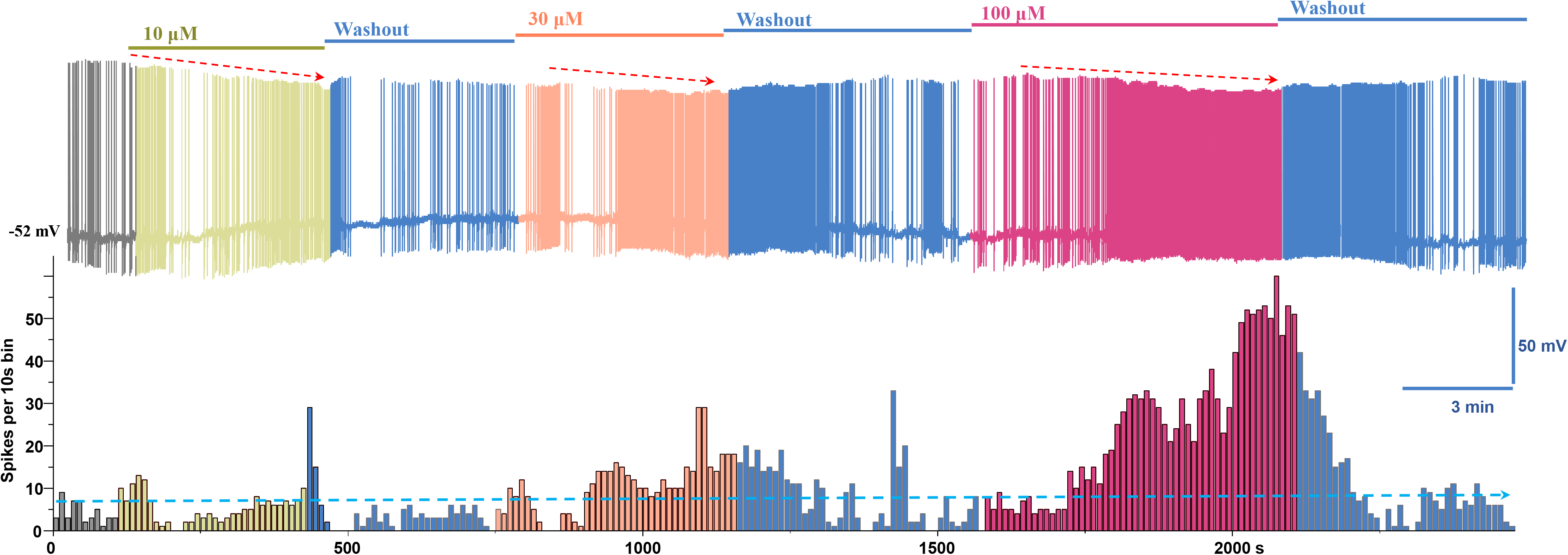
Manganese increases the spontaneous firing activity of dopamine neurons. Top: representative recording showing the dose-dependent effect of Mn^2+^ on the spontaneous firing activity of dopamine neurons and firing rate following washout period. Bottom: Histogram of firing frequency obtained from above trace.

**Figure 2.**
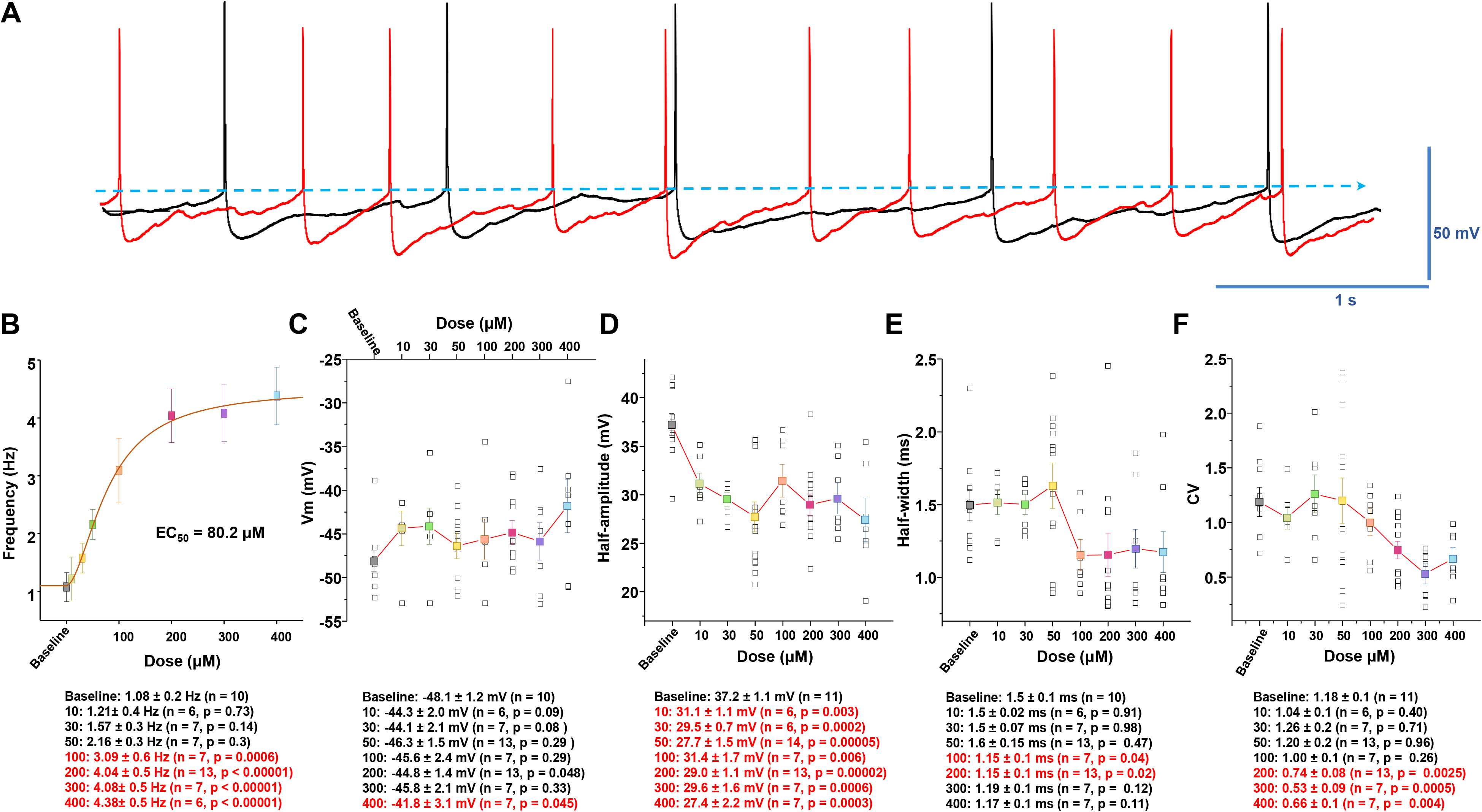
Analysis of spontaneous firing activity and properties of action potential following manganese exposure. ***A,*** Representative action potential trace before (baseline, black trace) and after bath application of 100 μM Mn^2+^ (red trace). ***B,*** Dose-response relationships of the spontaneous firing frequency of dopamine neurons following Mn^2+^ exposure. A EC_50_ of 80.2 μM was obtained by fitting the dose-dependent curve using a Hill equation. **C,** Mn^2+^ did not significantly change membrane potential. ***D,*** Mn^2+^ suppressed the amplitude of action potential (as measured as the half amplitude) at all concentrations examined (baseline: 37.2 ± 1.1 mV; 10 μM manganese: 31.1 ± 1.1 mV; *t*_(15)_ = 3.6, p = 0.03, two-tailed Student’s t tests; *n* = 11 vs 6). ***E,*** Mn^2+^ did not change the half-width of action potential at 10-50 μM, but it significantly narrowed the half-width at 100 μM (baseline: 1.5 ± 0.1 ms; 100 μM manganese: 1.15 ± 0.1 ms; *t*_(15)_ = 2.2, p = 0.04, two-tailed Student’s t tests; *n* = 10 vs. 7). ***F,*** Mn^2+^ dose-dependently decreased the coefficient-of-variation of interspike intervals (baseline: 1.2 ± 0.1; 200 μM manganese: 0.7 ± 0.1; *t*_(22)_ = 3.4, p = 0.003, two-tailed Student’s t tests; *n* = 11 vs 13).

Mice were injected intraperitoneally (i.p.) with manganese (II) chloride tetrahydrate (70 mg/kg, MnCl_2_·4H_2_O) in a sub-acute treatment based on a published protocol shown to significantly increase manganese concentrations in the basal ganglia (Dodd et al., 2005).

### Neuronal and cells culture

Neuronal primary culture was performed as previously described (Saha et al., 2014; Lin et al., 2016; Sambo et al., 2017). Briefly, mouse midbrain dopamine neurons from 0-2 day old pups of either sex were isolated and incubated in a dissociation medium (in mM): 116 NaCl; 5.4 KCl; 26 NaHCO_3_; 25 glucose; 2 NaH_2_PO_4_; 1 MgSO_4_; 1.3 cysteine; 0.5 EDTA; 0.5 kynurenate containing 20 units/mL papain at 34-36°C under continuous oxygenation for 2 hours. Tissue was then triturated with a fire-polished Pasteur pipette in glial medium (in %): 50 minimum essential media; 38.5 heat-inactivated fetal bovine serum; 7.7 penicillin/streptomycin; 2.9 D-glucose (45%) and 0.9 glutamine (200 mM). Dissociated cells were pelleted by centrifugation at 500 x g for 10 min and re-suspended in glial medium. Cells were plated on 12 mm round coverslips coated with 100 μg/ml poly-L-lysine and 5 μg/ml laminin in 35 x 10 mm tissue culture Petri dishes. One hour after plating, the medium was changed to neuronal medium: Neurobasal (Life Technologies, Grand Island, NY), 0.9% L-glutamine, 2% B27 and 1 ng/ml GDNF. Neuronal medium was conditioned overnight on cultured glia. The conditioned neuronal medium was supplemented with 1 ng/mL glial cell line-derived neurotrophic factor and 500 μM kynurenate and sterile-filtered before it was added to the neuronal cultures.

Plasmids for Ca_v_1.2-GFP and Ca_v_1.3-GFP were generous gifts from Dr. Gregory Hockerman (Purdue University). Briefly, cDNAs that encode the cytoplasmic II-III inter-domain loop of Ca_v_1.2 (amino acids 758-903) or Ca_v_1.3 (amino acids 752-885) were fused to enhanced GFP were constructed in the pEGFP-N1 vector (Clontech, Mountain View, CA). Plasmids were transformed into MAX Efficiency DH5α Competent Cells (Invitrogen, Waltham, MA) and amplified on Luria-Bertani (LB) agar plates with an ampicillin concentration of 50 mg/L. Individual colonies were selected and further amplified in 250 mL LB medium with 50 mg/L ampicillin. Plasmid DNAs were purified using the Qiagen Plasmid Plus Maxi Kit (Qiagen, Germantown, MD). The integrity of the clones was confirmed by cDNA sequencing and restriction digest analysis.

HEK 293 cells stably expressing Ca_v_1.2 or 1.3 were generous gift from Dr. Richard B. Silverman (University of Chicago). Cells were maintained in DMEM (ThermoFisher) supplemented with 10% fetal bovine serum (Gemini Cat. No. 100-106), 1.1 mM sodium pyruvate (Sigma-Aldrich Cat. No. S8636-100ML), 0.1 mg/mL Zeocin (InvivoGen Cat. No. ant-zn-1), and 0.05 mg/mL Hygromycin B (A.G. Scientific Cat. No. H-1012-PBS). Additionally, either 0.3 mg/mL G418 (Caisson Labs Cat. No. ABL06-20ML) or 0.2 mg/mL Blasticidin S hydrochloride (Research Products International Cat. No. B12200-0.05) was added for Ca_v_1.2 or Ca_v_1.3, respectively. Cells were passaged at 70-85% confluency 24-48 hours prior to experimentation using TrypLE Express Enzyme (Gibco Cat. No. 12604-021).

### ^54^Mn uptake

Effectene reagent (Qiagen, Valencia, CA) was used for transient transfection of HEK cells with a plasmid vector either empty or containing Ca_v_1.2 or Ca_v_1.3 cDNA. Overexpression of Ca_v_1.2 and Ca_v_1.3 was assessed by the western blotting 48 hours after transfection (Millipore Sigma). For ^54^Mn uptake experiments, following a 48-hour transfection, the cells were washed with Hanks’ balanced salt solution (HBSS) and incubated at 37°C in serum-free DMEM containing 40 µM MgCl_2_ and ^54^Mn at 0.18 µCi /mL (PerkinElmer). In some experiments, cells were pretreated with 10 µM Nifedipine 30 minutes before ^54^Mn treatment. After incubation with ^54^Mn for the times indicated, cells were washed with chelating buffer (10 mM EDTA, 10 mM HEPES, and 0.9% NaCl) and solubilized in 0.2% SDS 0.2 M NaOH for 1 hour. Radioactivity was measured by gamma-ray solid scintillation spectrometry. Protein content was measured colorimetrically with BCA reagent (Pierce-Thermo Fisher Scientific). Results were expressed as counts per minute (cpm) per mg total protein.

### Electrophysiological recordings

Spontaneous firing activity of midbrain dopamine neurons was examined via whole cell current clamp recordings as previously described (Saha et al., 2014; Lin et al., 2016; Sambo et al., 2017). The neurons were continuously perfused with artificial cerebral spinal fluid (aCSF) containing (in mM): 126 NaCl, 2.5 KCl, 2 CaCl_2_, 26 NaHCO_3_, 1.25 NaH_2_PO_4_, 2 MgSO_4_, and 10 dextrose, equilibrated with 95% O_2_-5% CO_2_; pH was adjusted to 7.4. Patch electrodes were fabricated from borosilicate glass (1.5 mm outer diameter; World Precision Instruments, Sarasota, FL) with the P-2000 puller (Sutter Instruments, Novato, CA). The tip resistance was in the range of 3-5 MΩ. The electrodes were filled with a pipette solution containing (in mM): 120 potassium-gluconate, 20 KCl, 2 MgCl_2_, 10 HEPES, 0.1 EGTA, 2 ATP, and 0.25 GTP, with pH adjusted to 7.25 with KOH. All experiments were performed at 37°C. To standardize action potential (AP) recordings, neurons were held at their resting membrane potential (see below) by DC application through the recording electrode. Action potential was recorded if the following criteria were met: a resting membrane potential of less than −35 mV and an action potential peak amplitude of >60 mV. Action potential half-width was measured as the spike width at the half-maximal voltage using Clampfit 10 software (Axon instruments, Foster City, CA). Steady-state basal activity was recorded for 2–3 min before bath application of the drug. For experiments involving drug application, each coverslip was used for only one recording. The spontaneous spike activity of midbrain dopamine neurons was obtained by averaging 1 min interval activities at baseline (before manganese) and after 7-10 min of manganese. The relationship between Mn concentration and neuronal firing frequency was fitted according to a Hill equation: *E = E_max_Mn^nH^/(EC_50_ + Mn^nH^)* where E is the predicted effect of Mn, E_max_ is the maximum effect, *n*H is the slope factor, and the *n*th root of EC_50_ gives an estimate of the midpoint of the activation curve.

### Recording from HEK Cells expressing GFP-α subunits

HEK293 cells stably expressing GFP-α subunits where generous gift from Dr. Robert Brenner (University of Texas Health Science Center at San Antonio). The cells were cultured as described previously (Goodwin et al., 2009; Saha et al., 2014; Wang et al., 2009). The cells were plated on glass coverslips (Electron Microscopy Sciences, Hatfield, PA). Green fluorescent protein expression was used to identify α subunits-expressing cells. Electrophysiology experiments were performed 3 days after plating the cells. Macropatch recordings were performed using the excised inside-out patch clamp configuration at 22 - 23°C. Patch pipettes were pulled to a final tip resistance of 1.5–3 MΩ and filled with the internal solution. The electrode solution was composed of the following (in mM): 20 HEPES, 140 KMeSO_3_, 2 KCl, and 2 MgCl_2_, pH 7.2. Bath solution was composed of a pH 7.2 solution of the following (in mM): 20 HEPES, 140 KMeSO_3_, and 2 KCl. Intracellular Ca^2+^ was buffered with 5 mM HEDTA to give the required 10 μM free [Ca^2+^] concentration calculated using the MAXCHELATOR program (Winmaxchelator software; Dr. Chris Patton, Stanford University, Pacific Grove, CA). Recordings and data acquisition were described in *Single-channel recordings above*. Mean current density was plotted against membrane potential (*I*–*V*) and was fitted by the Boltzmann equation: *I* = *I*_min_ + (*I*_max_ − *I*_min_)/(1 + exp − κ(*V* − *V*_1/2_)), where *I* is the current, *I*_max_ is the maximum current, *I*_min_ is the minimum current, κ is a slope factor, and *V*_1/2_ is the midpoint potential.

### Pharmacological isolation of Ca^2+^ currents

For whole-cell recordings, we used a Cesium methanesulfonate-based intracellular solution composited in mM: 110 CsMeSO_3_, 10 TEA-Cl, 10 4-aminopyridine (4-AP), 10 HEPES, 1 MgCl_2_, 10 EGTA, 2 ATP and 0.3 GTP, with pH adjusted to 7.25 with CsOH. Bath solution contained following (in mM): 110 NaCl, 35 tetraethylammonium (TEA)-Cl, 1 CsCl, 1 MgSO_4_, 2 CaCl_2_, 10 HEPES, 11.1 dextrose and 0.001 tetrodotoxin (TTX), pH 7.4 with CsOH. For the Ca^2+^-free bath solution, 2 mM CaCl_2_ was replaced with 3 mM MgCl_2_, a HEPES solution containing 4 mM Mg^2+^. Calcium currents were evoked by a series of 200-ms depolarizing steps from −60 to +85 mV in 5-mV increments. To avoid possible interference between responses, the depolarizing voltage steps were delivered every 5 s. Data were obtained in 1- to 3-min intervals while the patches were held at −65 mV. The series resistances were in the range of 5–10 MΩ (typically 5 MΩ) and were compensated 60% on-line. Membrane potential measurements were not corrected for the liquid junction potential (∼15 mV). Leak currents were subtracted using a standard P/4 protocol. Before seals (5 GΩ) were made on cells, offset potentials were nulled. Capacitance subtraction was used in all recordings. To determine current density (pA / pF), the peak current value of the steady-state current at 180 ms was divided by the membrane capacitance from each recorded cell.

### Preparation of mouse brain slices

Unless otherwise noted, 30-40 day-old C57BL/6 male mice in either wild type or mice conditionally expressing GCaMP6f in dopaminergic neurons (described above) were used for slice preparation. Only males heterozygous were selected and used for these experiments. Mice were deeply anesthetized with 4% isoflurane, and the brain was subsequently removed from the cranium. The tissue was glued onto the cutting stage and submersed in ice-cold, oxygenated aCSF (equilibrated with 95% O_2_-5% CO_2_). Coronal or horizontal brain slices (200 μm) containing the substantia nigra compacta or striatum were cut using a MicroSlicer Zero 1N (Dosaka, Kyoto, Japan).

### FFN200 loading and multiphoton imaging

Striatal slices of wild type mice were incubated in oxygenated aCSF with 10 μM FFN200 (Tocris, Minneapolis, MN) for 30 min at 22–24° C and washed for 45–50 min before imaging. After incubation with FFN200, dorsal striatal slices were transferred into imaging chambers, then continuously perfused with aCSF equilibrated with 95% O_2_-5% CO_2_. *In vitro* multiphoton imaging was performed with a Nikon AR1 MP (Nikon Instruments Inc., Melville, NY, USA) equipped with a plan apo LWD 25x, water-immersion (NA=1.1, WD=1.43 - 2.04mm, DIC N2) objective. Multiphoton excitation was generated with a Spectra Physics 15 W Mai Tai eHP tunable Ti:sapphire femtosecond pulsed IR laser. FFN200 imaging experiments were performed using excitation wavelengths of 830-840 nm and an emission of 430–470 nm. Emission light was directed with a 405 nm dichroic mirror through a 340 nm short pass filter to enhanced hybrid PMTs via fluorophore specific filters. Imaging sites in time lapse experiments were aligned based on the multiphoton imaged dendritic structure and FFN200 puncta. Imaging sites in time lapse experiments were aligned based on the multiphoton imaged dendritic structure and FFN200 puncta. Images were collected with a 2.4 μs pixel dwell time, with the laser attenuated to 3% of total power, a pixel size of 0.24 μm (XY), and a step size of 0.5 μm (Z) at 1,024 × 1,024 pixel resolution. After acquiring an initial image (t = 0), perfusion was switched to aCSF containing 100 μM MnCl_2_ and images acquired every 4s. Control slices labeled with FFN200 were imaged 5 – 10 min in the absence of MnCl_2_.

Two photon images and analysis of fluorescent intensity for FFN200 puncta were performed using NIS-Elements Software v4.5 (Nikon Instruments Inc., Melville, NY, USA). Regions of interest (ROIs) were automatically selected by the program. Background fluorescence intensity was determined as the mean fluorescence intensity of areas of the image that excluded puncta. The mean background-subtracted fluorescent puncta intensity was then calculated by subtracting the mean background intensity from the mean fluorescence intensity measured at the puncta.

### Fura-2 calcium imaging

For Fura-2 calcium imaging, primary cultured midbrain neurons (8-10 DIV) were used in all studies. Neurons were washed twice in HBSS (Life Technologies) followed by 30-min treatment in 5 μM Fura-2 AM (Thermo Fisher). Cells were then washed twice with HBSS followed by 30-min incubation in HBSS. Cells were then imaged on a Nikon Ti Eclipse inverted microscope. Fluorescence was monitored using dual excitation wavelengths (340/380 nm) and a single emission wavelength (510 nM). Fura-2 bound to free calcium is excited at 340 nm, whereas unbound Fura-2 is excited at 380 nm. Baseline images were taken for 30 s to 1 min followed by addition of vehicle, 100 μM manganese, or 20 mM caffeine as the positive control group. The concentration of caffeine used in this study was selected based on a previous study examining caffeine-regulation of Ca^2+^ mobilization in the midbrain dopamine neurons (Choi et al., 2006). Images were acquired for 4 min after treatment. For analysis, experimenter defined regions of interest (ROIs) were drawn along the cell body of each neuron. Ca^2+^ changes were determined as the fluorescence normalized to the average fluorescence of the first 30 s of baseline imaging for each neuron. The percent fluorescence change over baseline was calculated as: %_Δ_F/F_0_ = (F_max_-F_0_)/F_0_ x 100, where F_max_ is the maximal fluorescent value after treatment and F_0_ is the average fluorescence at baseline. Images were presented as the ratio of 340/380 imaging.

### Total internal reflection fluorescence microscopy (TIRFM)

For these experiments, HEK293 cells expressing GFP-α subunits were plated onto 35 mm glass-bottom dishes (No. 1 thickness) to 60-80% confluence. Nikon Ti Eclipse (Nikon, Melville, NY) inverted microscope equipped with a multi-line solid-state laser source (470, 514, 561nm) was used for all TIRFM imaging. Lasers were guided through a 60X 1.4 NA objective. Images were detected digitally using a CCD camera. Image exposure time was coupled with stimulation duration at 100 ms and laser intensity was maintained at 40%. For quantification of fluorescence intensity at the cell surface, experimenter-defined ROIs were created for each cell in order to exclude both cell membrane overlap between adjacent cells and measurement of intensity at the peripheral edges of cells. Background fluorescence was subtracted from all images. Mean intensity over time for each ROI was recorded continuously before and after the application of 100 µM manganese to the bath solution. Experiments were performed in the isotonic, isosmotic external solution described above. The baseline fluorescent intensity is defined as the average fluorescent intensity 1 min prior to drug application. All values were normalized to the baseline fluorescent intensity. Single count elementary sequential smoothing was applied for image presentation only, and not for analysis. Bleaching at each experimental time point was determined by the amount of decreased fluorescence intensity in untreated cells.

### Magnetic Resonance Imaging Scans

Seventeen adult male mice (35-40 days-old male wild-type C57BL/6 mice) were randomly assigned to two groups either receiving manganese only or manganese plus nifedipine. Systemic injections of L-type Ca^2+^ channel inhibitor, nifedipine (15 mg/kg), which can effectively cross the blood–brain barrier (Cain et al., 2002), were used to inhibit the L-type Ca^2+^ channels (Jinnah et al., 1999). The treatment doses were based on the previously published protocols (Cain et al., 2002, Dodd et al., 2005, Jinnah et al., 1999). Both groups were intraperitoneal (i.p.) injected with manganese (II) chloride tetrahydrate (70 mg/kg), but one group received manganese injection after 30 min nifedipine (15 mg/kg, i.p) injection. Magnetic resonance (MR) scanning was performed twenty-four hours after manganese exposure. On the scanning day, mice were induced using 3-4% isoflurane delivered in medical grade air (70% nitrogen, 30% oxygen; air flow rate 1.5 mL/min). The anesthesia was maintained at 1.0-1.5% isoflurane during MRI scanning. Core body temperature and spontaneous respiratory rates were continuously recorded during MRI scanning (SA Instruments, Stony Brook, NY). Mice were maintained at normal body temperature levels (37– 38 °C) using a warm water recirculation system. The MEMRI scans were collected in a 4.7T/33 cm horizontal bore magnet (Magnex Scientific) at the Advanced Magnetic Resonance Imaging and Spectroscopy facility in the McKnight Brain Institute of the University of Florida. The MR scanner consisted of a 11.5 cm diameter gradient insert (Resonance Research, Billerica, MA, USA) controlled by a VnmrJ 3.1 software console (Agilent, Palo Alto, CA, USA). A quadrature transmit/receive radiofrequency (RF) coil tuned to 200.6 MHz ^1^H resonance was used for B1 field excitation and RF signal detection (airmri; LLC, Holden, MA). The MEMRI included a multiple repetition time sequence to calculate parametric T_1_ maps for each group using a fast spin echo sequence with a total of four TR’s (0.5, 1.08, 2.33, 5.04 seconds), and TE = 6.02 ms with the following geometric parameters: 16 x 16 mm^2^ in plane, 14 slices at 0.8 mm thickness per slice, data matrix = 128 x 128 (125 μm in-plane resolution).

### MRI Post-Processing

Whole brain masks were obtained via automatic segmentation with PCNN using high-resolution anatomical scans to remove non-brain voxels. All cropped data were used to create templates for each cohort using Advanced Normalization Tools (ANTs; http://stnava.github.io/ANTs/). The templates were then registered to an atlas of the mice brain using the FMRIB Software Library linear registration program flirt (Jenkinson et al., 2002). The atlas was then transformed back to each individual data set with the registration matrices from ANTS. To generate parametric T1 maps, multi-TR images were fit to the equation S_TR_=S_0_(1-e^-TR/T1^) using non-linear regression in QuickVol II for ImageJ (Schmidt et al., 2004). From T1 maps, the T1 relaxation rate (R1 in ms^-1^) is calculated and exported from regions of interest. These methods allowed the measurement of the amount of activity, along with the location of manganese uptake in both the control and drug exposed groups.

### Data acquisition and analysis

The reported “n” for each experiment is obtained from at least three independent biological replicates. All statistical analyses were performed on data presented in the manuscript are stated in the figure legends and detailed in the methods and the results section. For each experiment, statistical tests were chosen based on the structure of the experiment and data set. No outliers were removed during statistical analysis. Sample sizes estimates were based on published biological, biochemical and electrophysiological literature that utilized the animal model; this was within a range commonly employed by researchers in our field using similar techniques. The electrophysiology data were acquired using the ClampEx 10 software (Molecular Devices). The data were analyzed off-line using pClamp 10. For all experiments, the data are presented as mean ± SEM. N denotes the number of neurons, cells, or mice for each experiment. Statistical significance was assessed using two-tailed Student’s *t* tests or one-way ANOVA. If ANOVA showed statistical significance, all pairwise *post hoc* analysis was performed using a Tukey’s *post hoc* test. Differences were considered significant at *P* < 0.05. * denotes significance *p* < 0.05. ** denotes significance *p* < 0.01. The coefficient of variation is a measure of the relative spread of the data. It is computed as the standard deviation divided by the mean times 100%. SigmaPlot 11 was used for all statistical analysis.

## Results

We exploited multiple complementary approaches to identify dopamine neurons for the electrophysiological recordings, as described previously (Lin et al., 2016). A total of 149 midbrain dopamine neurons were recorded and analyzed as outlined below. The primary parameters of passive membrane and action potential were averaged in 30 randomly selected cells. The average resting membrane potential was −48.3 ± 1.1 mV; input resistance was 252.5 ± 21.9 MΩ; membrane time constant was 826.9 ± 52.5 μs; and membrane capacitance was 64.1 ± 3.3 pF.

### Manganese altered the intrinsic firing behavior of midbrain dopaminergic neurons

Spontaneous firing activity of dopamine neurons was measured before and after manganese application at concentrations of 10, 30, and 100 μM (Figure 1). Lower concentrations of 10 and 30 μM are within the normal range of intracellular manganese (Bowman and Aschner, 2014), whereas 100 μM falls within the pathological range (Gandhi et al., 2018). Acute exposure to manganese increased the spontaneous firing frequency and suppressed the amplitude of action potentials (AP) (red arrows) in a concentration-dependent manner. Superimposed representative traces of spontaneous spikes before (black trace) and after 100 μM manganese (red trace) exposure revealed that manganese increased the firing frequency of dopamine neurons and truncated AP amplitudes (Figure 2A). The EC_50_ for the manganese-mediated increase in firing frequency of dopamine neurons was determined to be 80.2 μM by fitting a concentration-dependent curve using a Hill equation (Figure 2B). While manganese induced a modest membrane depolarization at lower concentrations (P > 0.05), it significantly depolarized membrane potential at the highest concentration tested in this study (100 μM), which is consistent with the reported manganese concentration for *in vitro* studies (Bowman and Aschner, 2014) (Figure 2C; baseline: −48.1 ± 1.2 mV; 400 μM manganese: −41.8 ± 3.1 mV; *t*_(15)_ = −2.2, p = 0.045, two-tailed Student’s t tests; *n* = 7 vs 10). Although manganese markedly suppressed AP amplitude at the lowest dose of 10 μM (baseline: 37.2 ± 1.1 mV; 10 μM manganese: 31.1 ± 1.1 mV; *t*_(15)_ = 3.6, p = 0.03, two-tailed Student’s t tests; *n* = 11 vs 6), higher doses of manganese did not further suppress AP amplitude (Figure 2D). In contrast, manganese did not influence AP half-width at low concentrations, but it significantly narrowed AP width at 100 μM (Figure 2E, baseline: 1.5 ± 0.1 ms; 100 μM manganese: 1.15 ± 0.1 ms; *t*_(15)_ = 2.2, p = 0.04, two-tailed Student’s t tests; *n* = 10 vs. 7). Manganese progressively decreased the coefficient-of-variation (CV) from 50 μM, indicating a reduction in firing variation evident by the small interspike interval (Figure 2F, baseline: 1.2 ± 0.1; 200 μM manganese: 0.7 ± 0.1 ms; *t*_(22)_ = 3.4, p = 0.003, two-tailed Student’s t tests; *n* = 11 vs 13). These results suggest manganese can regulate the intrinsic firing behavior of dopamine neurons at both physiological and pathophysiological concentrations in a concentration-dependent manner. To address the pathological effects of manganese, 100 μM was used for the rest of the study based on the determined EC_50_ in Figure 2B.

### Manganese-stimulation of neuronal activity increased the release of fluorescent false neurotransmitter (FFN200)

Next, we asked whether manganese-stimulation of firing frequency of dopamine neurons leads to increased neurotransmitter release. To do this, we utilized FFN200, a fluorescent substrate of VMAT2 that selectively traces monoamine exocytosis in both neuronal culture and striatal slices (Pereira et al., 2016). “De-staining” of FFN200 associated with monoamine vesicular release has been used as a surrogate for neurotransmitter release (Pereira et al., 2016). Consistent with a previous report (Pereira et al., 2016), we found following vehicle (aCSF) application there is a time-dependent and slow de-staining of FFN200 at striatal dopamine terminals (Supplemental Figure 1), suggesting a direct correlation between spontaneous firing activity of dopamine neurons and neurotransmitter release. Manganese increased FFN200 release as measured by increased rate and magnitude of fluorescence de-staining (see supplemental movie 1). These data suggest manganese-stimulation of neuronal activity correlates with increased neurotransmitter release (see yellow arrows in Supplemental Figure 1, mean ± SEM, *F*_(1,148)_ = 70.2, †: *p* < 0.01, one-way ANOVA followed by Tukey’s test, *n* = 4 slices / group). Additional control experiments were performed when the release of FFN200 is inhibited. Since blockade of dopamine transporter (DAT) decreases firing activity of dopamine neurons (Saha et al., 2014; Lin et al., 2016), theoretically, it should decrease the de-staining rate or even increase the fluorescent punctate due to accumulation of FFN200-filled synaptic vesicles in the terminal regions. As shown in Supplemental Figure 1, treatment with DAT inhibitor nomifensine (5 μM) produced a plateau in the fluorescent signal followed by increased fluorescent levels, suggesting inhibition of neurotransmitter release. These data further support the findings that manganese treatment increases the activity of dopaminergic neurons.

### Removal of extracellular Ca^2+^ did not prevent manganese stimulation of spontaneous spike frequency

Our observed broad action potentials and slow autonomous pacemaking activities are consistent with typical electrophysiological features in dopamine neurons (Bean, 2007). Numerous studies have shown that slow rhythmic spiking is accompanied by large oscillations in intracellular Ca^2+^ concentrations that are driven by the opening of voltage-dependent Ca^2+^ channels (Guzman et al., 2009; Mercuri et al., 1994; Nedergaard et al., 1993; Puopolo et al., 2007). Therefore, next we measured manganese-regulation of the spontaneous firing activity of dopamine neuron in Ca^2+^– free extracellular solution. Unexpectedly, removal of extracellular Ca^2+^ did not prevent manganese-stimulation of the firing frequency (Figure 3A, C; baseline: 1.3 ± 0.2 Hz vs. manganese in Ca^2+^–free: 6.5 ± 1.1 Hz; *F*_(2,18)_ = 20.9, *p* = 0.000002, one-way ANOVA followed by Tukey’s test; *n* = 7 / group). Ca^2+^–free solution containing manganese did not change the membrane potential (Figure 3D), AP half-width (Figure 3E), or CV (Figure 3G); however, it significantly truncated the amplitude of AP (Figure 3F; baseline: 45.4 ± 2.2 mV vs. manganese in Ca^2+^–free: 37.9 ± 0.7 mV; *F*_(2,18)_ = 16.1, *p* = 0.0001, one-way ANOVA followed by Tukey’s test; *n* = 7 / group). In the absence of manganese, comparison of the firing activity before and after perfusion of a Ca^2+^–free external solution revealed a reduction in the firing frequency (Figure 3C, 0.5 ± 0.03 Hz), broadening of AP half-width (Figure 3E: baseline: 1.6 ± 0.08 ms vs Ca^2+^–free: 2.9 ± 0.2 ms, *F*_(2,18)_ = 35.9, *p* = 0.0001, one-way ANOVA followed by Tukey’s test; *n* = 7 / group), truncated AP amplitude (Figure 3F, 32.5 ± 2.0 mV, *n* = 7 / group), and an increased CV (Figure 4G, baseline: 1.1 ± 0.2 vs Ca^2+^–free: 1.99 ± 0.3, *F*_(2,18)_ = 7.3, *p* = 0.004, one-way ANOVA followed by Tukey’s test; *n* = 7 / group). These results suggest that manganese stimulation of neuronal activity it is not dependent on extracellular Ca^2+^ influx into the dopamine neuron.

**Figure 3.**
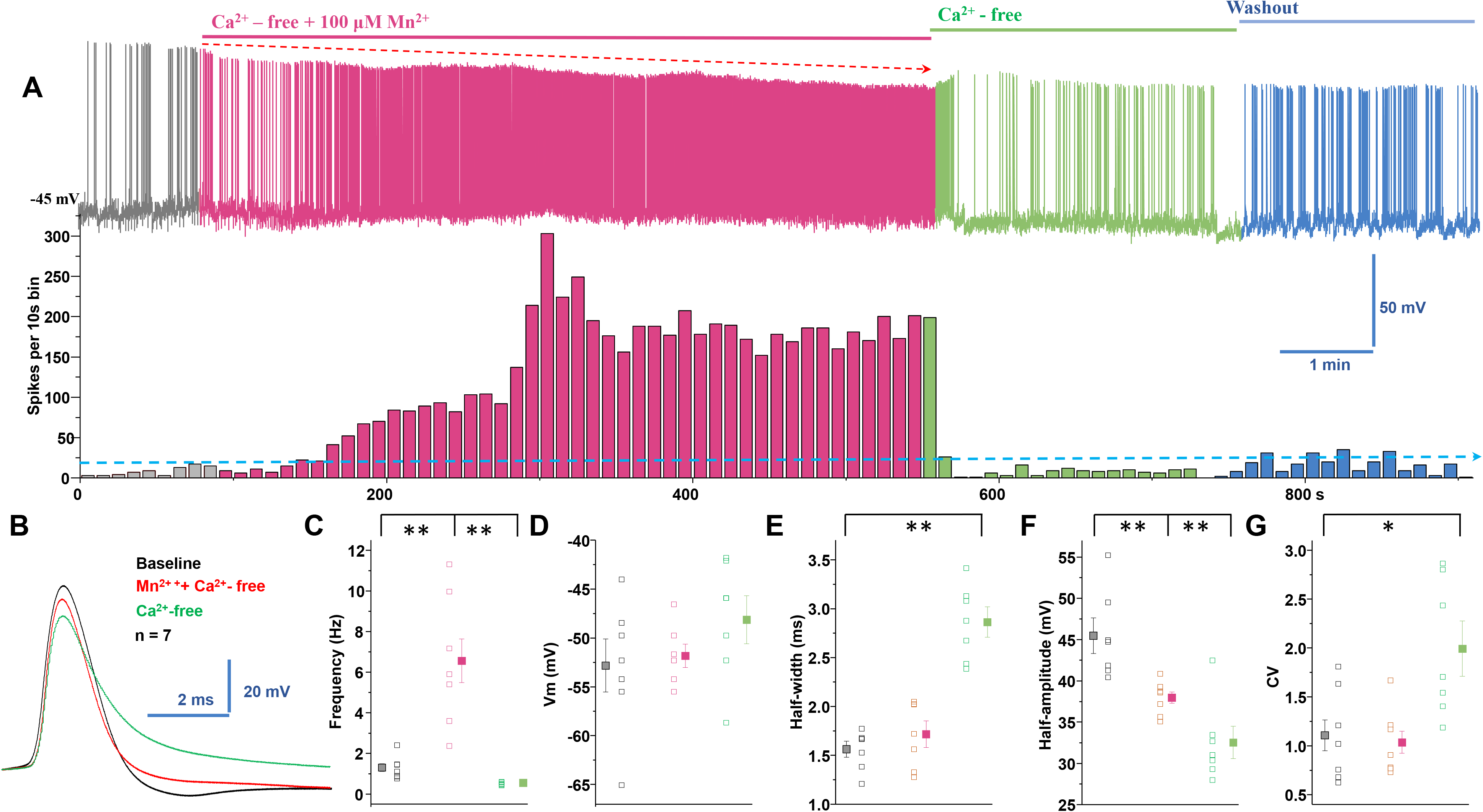
Manganese stimulation of firing activity of dopamine neurons is not dependent on extracellular Ca^2+^. ***A,*** Top: representative spontaneous spike activities of a neuron exposed to Mn^2+^ in a Ca^2+^-free extracellular solution. Middle: histogram of firing frequency obtained from the above trace showing Mn^2+^ increased firing frequency. ***B,*** Superimposed representative single action potential traces shown in A, baseline (black trace), Mn^2+^ + Ca^2+^-free solution (red trace), and Ca^2+^-free solution only (green trace). ***C,*** In Ca^2+^-free solution, Mn^2+^ increased spontaneous firing rate. Mn^2+^ washout in Ca^2+^-free conditions revealed a reduction of firing rate (baseline: 1.3 ± 0.2 Hz vs. manganese in Ca^2+^ – free: 6.5 ± 1.1 Hz; *F*_(2,18)_ = 20.9, *p* = 0.000002, one-way ANOVA followed by Tukey’s test; *n* = 7 / group). ***D,*** Mn^2+^ and Ca^2+^-free extracellular solution did not change membrane potential. ***E,*** Half-width measured at half-maximal voltage of action potential was not different between baseline and Mn^2+^ application, but it was significantly broadened in Ca^2+^-free condition as compared to baseline (baseline: 1.6 ± 0.08 ms vs Ca^2+^ – free: 2.9 ± 0.2 ms, *F*_(2,18)_ = 35.9, *p* = 0.0001, one-way ANOVA followed by Tukey’s test; *n* = 7 / group). ***F,*** Both Ca^2+^-free only and Mn^2+^ plus Ca^2+^-free condition significantly depressed the amplitude of action potential (baseline: 45.4 ± 2.2 mV vs. manganese in Ca^2+^ – free: 37.9 ± 0.7 mV; *F*_(2,18)_ = 16.1, *p* = 0.0001, one-way ANOVA followed by Tukey’s test; *n* = 7 / group). ***G,*** The coefficients of variation of the interspike intervals was not different between baseline and Mn plus Ca^2+^-free solution (baseline: 1.1 ± 0.2 vs Ca^2+^ – free: 1.99 ± 0.3, *F*_(2,18)_ = 7.3, *p* = 0.004, one-way ANOVA followed by Tukey’s test; *n* = 7 / group). *p < 0.05; **p < 0.01.

**Figure 4.**
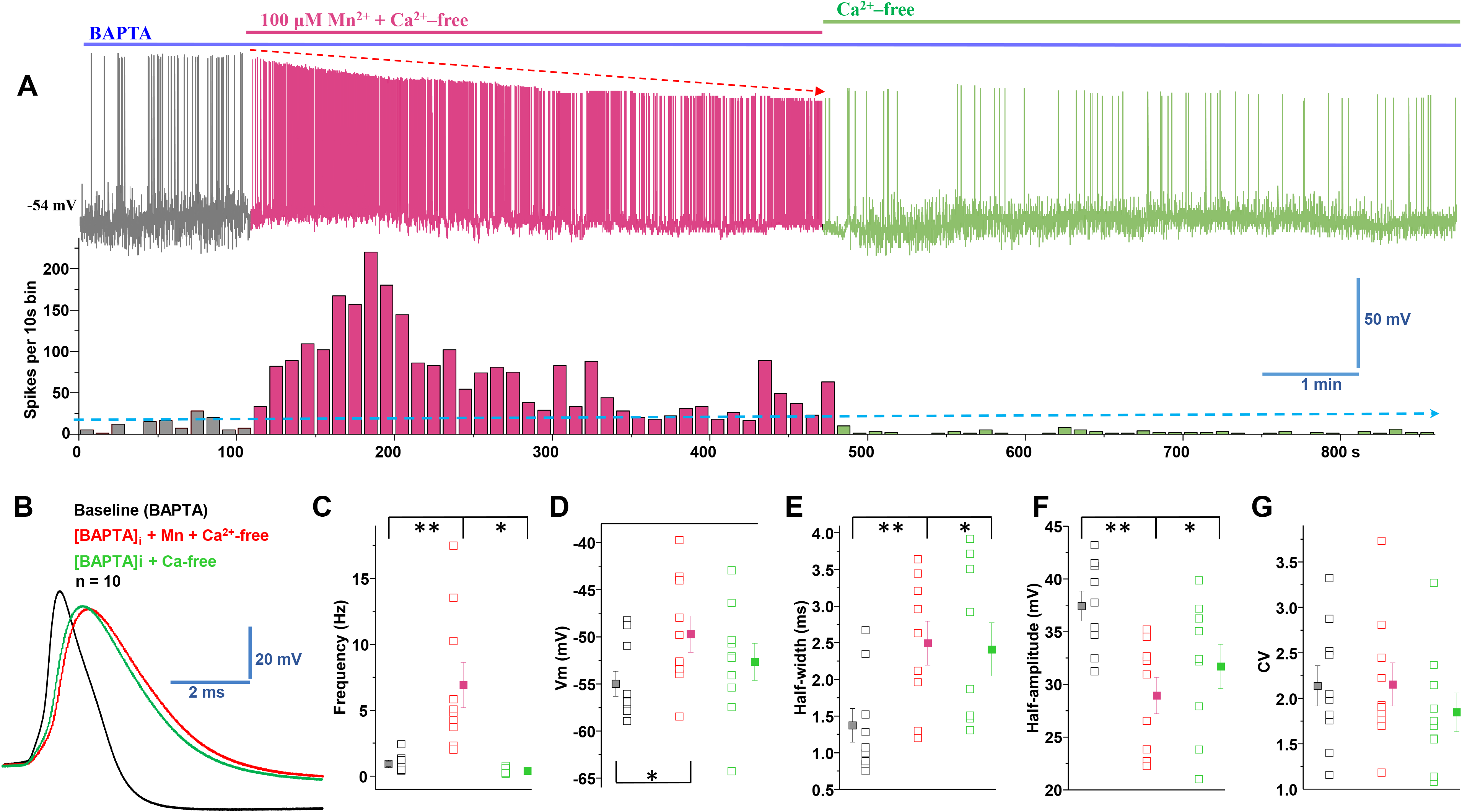
Manganese increases firing rate in the combined absence of extracellular and intracellular Ca^2^. ***A,*** Top: representative trace of spontaneous spike activity following Mn^2+^ exposure in an Ca^2+^-free extracellular solution and an intracellular solution containing the Ca^2+^ chelator BAPTA (10 mM). Bottom: histogram of firing frequency obtained from above trace showing Mn^2+^ increased firing frequency. ***B,*** Superimposed representative single action potential traces are shown in A: baseline (black trace), 100 μM Mn^2+^ + Ca^2+^-free + BAPTA (red trace) and Ca^2+^-free plus BAPTA-AM (green trace). ***C,*** Mn^2+^ increased firing rate even in the combined absence of extracellular and intracellular Ca^2+^; after Mn^2+^ washout, the firing rate decreased under Ca^2+^-free conditions (baseline: 0.9 ± 0.2 Hz vs. 100 μM manganese: 6.9 ± 1.7 Hz, *F*_(2,26)_ = 13.2, *p* = 0.003, one-way ANOVA followed by Tukey’s test; *n* = 10 / group). ***D,*** The combination of extracellular Ca^2+^-free plus Mn^2+^ and intracellular BAPTA containing solution depolarized the membrane potential. After Mn^2+^ washout the membrane potential returned to the baseline level (Ca^2+^-free conditions) (baseline: −54.9 ± 1.3 mV vs. 100 μM manganese: −49.6 ± 1.9 mV; *F*_(2,27)_ = 2.5, *p* = 0.028, one-way ANOVA followed by Tukey’s test; *n* = 9 / group). ***E,*** The half-width was measured at the half-maximal voltage of APs. The half-width was broadened significantly in the combination of extracellular Ca^2+^-free + intracellular BAPTA containing solution in the absence or presence of Mn^2+^ (baseline: 1.37 ± 0.2 ms vs. 100 μM manganese: 2.50 ± 0.3 ms; *F*_(2,24)_ = 3.5, *p* = 0.009, one-way ANOVA followed by Tukey’s test; *n* = 10 / group). ***F,*** Half-amplitude of action potential was significantly depressed in the combination of extracellular Ca^2+^-free and intracellular BAPTA contained solution in the absence or presence of Mn^2+^ (baseline: 37.5 ± 1.4 mV vs. 100 μM manganese: 29.0 ± 1.7 mV; *F*_(2,24)_ = 6.0, *p* = 0.001, one-way ANOVA followed by Tukey’s test; *n* = 9 / group). ***G,*** The coefficients of variation of the interspike intervals were not significantly different amongst the experimental groups. *p < 0.05; **p < 0.01, n = 9 - 10 per group.

### Combined application of Ca^2+^–free external solution and chelation of intracellular Ca^2+^ did not prevent manganese-stimulation of firing frequency

Since excluding extracellular Ca^2+^ (Ca^2+^**–**free external solution) did not decrease the effect of manganese on the spontaneous firing activity of dopamine neurons, AP half-width, amplitude, and CV (Figure 3), we asked whether manganese promotes neuronal activation by increasing intracellular Ca^2+^ concentrations([Ca^2+^]_i_) (Tjalkens et al., 2006). To test this, neurons were pretreated with BAPTA-AM (10 mM), a membrane permeable Ca^2+^ chelator, to deplete [Ca^2+^]_i_ followed by application of manganese in the presence of Ca^2+^–free external solution. Consistent with previous reports (Benedetti et al., 2011; Torkkeli et al., 2012), BAPTA pretreatment prior to bath application of manganese in Ca^2+^**–**free external solution (denoted in blue) depolarized the membrane potential (Figure 4D, baseline: - 54.9 ± 1.3 mV vs. 100 μM manganese: −49.6 ± 1.9 mV; *F*_(2,27)_ = 2.5, *p* = 0.028, one-way ANOVA followed by Tukey’s test; *n* = 9 / group) and broadened the AP half-width (Figure 4B, E baseline: 1.37 ± 0.2 ms vs. 100 μM manganese: 2.50 ± 0.3 ms; *F*_(2,24)_ = 3.5, *p* = 0.009, one-way ANOVA followed by Tukey’s test; *n* = 10 / group). Interestingly, we found that manganese treatment in Ca^2+^**–**free external solution (denoted in red) still increased the spontaneous firing activity of the neurons (Figure 4A, C, baseline: 0.9 ± 0.2 Hz vs. 100 μM manganese: 6.9 ± 1.7 Hz, *F*_(2,26)_ = 13.2, *p* = 0.003, one-way ANOVA followed by Tukey’s test; *n* = 10 / group) and truncated the amplitude of APs (Figure 4B, F; baseline: 37.5 ± 1.4 mV vs. 100 μM manganese: 29.0 ± 1.7 mV; *F*_(2,24)_ = 6.0, *p* = 0.001, one-way ANOVA followed by Tukey’s test; *n* = 9 / group). Washout of extracellular manganese in neurons pretreated with BAPTA and recorded in Ca^2+^**–**free external solution (denoted in green) exhibited a significant decrease in the firing frequency (Figure 4C, 0.4 ± 0.08 Hz), a broadening of AP half-width (Figure 4B, E:2.4 ± 0.4 ms, F_(2,24)_ = 4.3, p = 0.029, one-way ANOVA followed by Tukey’s test; n = 9 / group), and truncated AP amplitude (Figure 4B, F, 31.7 ± 2.1 mV, F_(2,24)_ = 6.0, p = 0.028, one-way ANOVA followed by Tukey’s test; n = 9 / group). These results suggest manganese-enhancement of firing frequency of dopamine neurons (Figure 1-3) is not due to the release of Ca^2+^ from intracellular stores.

### Manganese does not affect intracellular Ca^2+^ homeostasis in midbrain dopamine neurons

To directly examine the effect of manganese exposure on [Ca^2+^]_i_ in dopamine neurons, we measured Ca^2+^ responses in the midbrain slices from mice conditionally expressing GCaMP6f in dopaminergic neurons. Brain slices were continuously perfused with aCSF containing manganese or N-Methyl-d-aspartate (NMDA) equilibrated with 95% O_2_-5% CO_2_. Since NMDA receptors are Ca^2+^-permeable glutamate receptors (Choi, 1987; Wild et al., 2014), 100 μM NMDA was used as a positive control. Representative two-photon Ca^2+^ images are shown in Figure 5, where images for manganese (Figure 5A) or NMDA (Figure 5B) were taken for 10 min after 10 min of baseline imaging. We found manganese exposure (100 μM) did not significantly alter [Ca^2+^]_i_ compared to vehicle control. In contrast, 100 μM NMDA markedly increased [Ca^2+^]_i_ in the neurons. The relative intensity of two-photon Ca^2+^ signaling over baseline over time is shown in Figure 5C, and supplemental movie 2, and 3. The signal intensity was not affected by manganese administration (black) [Ca^2+^]_i_ whereas NMDA (red) robustly increased the GCaMP6f response, especially in neurons with low baseline fluorescence (*F*_(1,592)_ = 115.1, †: *p* < 0.001, one-way ANOVA followed by Tukey’s test, *n*: Mn = 10; NMDA = 11). In complement, ratiometric Ca^2+^ imaging using FURA-2 AM in cultured dopamine neurons confirmed the findings that manganese does not affect intracellular Ca^2+^ homeostasis in primary culture midbrain dopamine neurons (Supplemental Figure 2). Collectively, the lack of effect of manganese on Ca^2+^ homeostasis both in midbrain slices and culture is consistent with the notion that the effects of manganese on dopamine neurons are not a result of Ca^2+^ modulation.

**Figure 5.**
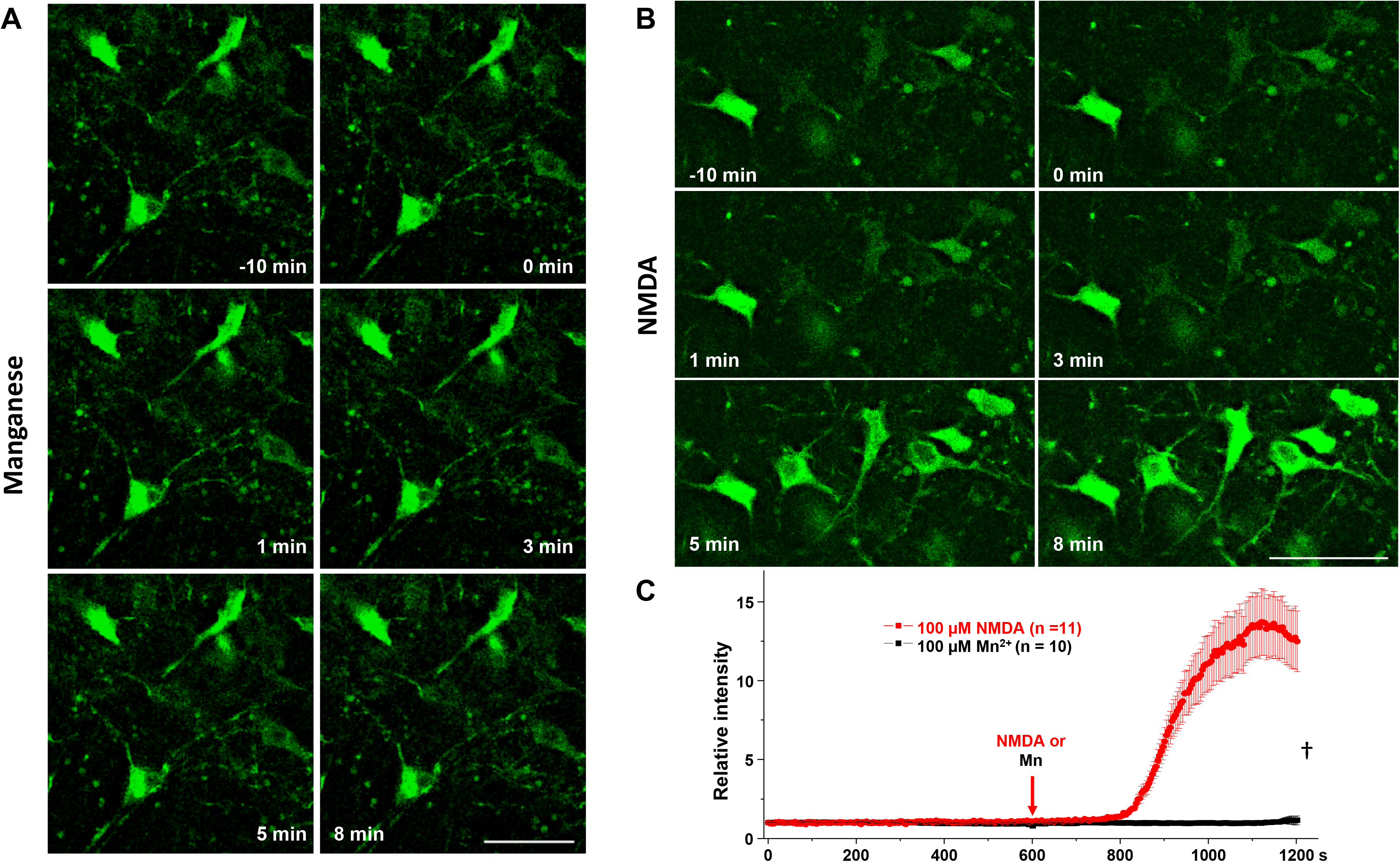
Manganese does not alter intracellular Ca^2+^ homeostasis. ***A, B,*** Representative two-photon images of GCaMP6f fluorescent neurons in mouse midbrain slices following 100 μM Mn^2+^ or NMDA treatments. ***C,*** Analyses of relative fluorescence intensities of GCaMP6 neurons following 100 μM Mn^2+^ or NMDA (*F*_(1,592)_ = 115.1, †: *p* < 0.001, one-way ANOVA followed by Tukey’s test, *n*: Mn = 10 frame (71 neurons) NMDA = 11 frame (50 neurons) from 3 independent experiments). Scale bar: 50 μm. †: p < 0.001.

### Cadmium blockade of voltage-gated Ca^2+^ channels inhibited manganese-stimulation of spontaneous firing activity of dopamine neurons

Thus far, above data suggest manganese enhances the spontaneous activity of midbrain dopamine neurons, but rhythmic spiking changes during administration of manganese are unlikely to involve extracellular Ca^2+^ influx or cytosolic Ca^2+^ release from intracellular stores. To further corroborate whether or not manganese augments the spontaneous activity of dopamine neurons through Ca^2+^ channels, we examined the effect of a nonselective Ca^2+^ channel blocker, cadmium (Cd^2+^, 100 μM) on the intrinsic firing behaviors of dopamine neurons following manganese exposure. As shown in Figure 6, nonselective blockade of Ca^2+^ channels by bath application of Cd^2+^ suppressed the spontaneous firing rate of dopamine neurons (Figure 6A). Unexpectedly, we found under this condition, manganese application failed to enhance the firing frequency (Figure 6A, C: *F*_(2,18)_ = 8.3, *p* = 0.002, one-way ANOVA followed by Tukey’s test, *n* = 7 group). Treatment with Cd^2+^ alone or the co-administration of Cd^2+^ and manganese did not change the membrane potential (Figure 6D), AP half-width (Figure 6B, E), and AP half-amplitude (Figure 6B, F), but as expected significantly increased the CV (Figure 6G: *F*_(2,18)_ = 4.6, *p* = 0.024, one-way ANOVA followed by Tukey’s test, *n* = 7 group). These findings suggest that despite observations that manganese-mediated increase in firing activity of dopamine neurons is not dependent on extracellular or intracellular Ca^2+^ flux, it is dependent on Ca^2+^ channels expressed on dopamine neurons.

**Figure 6.**
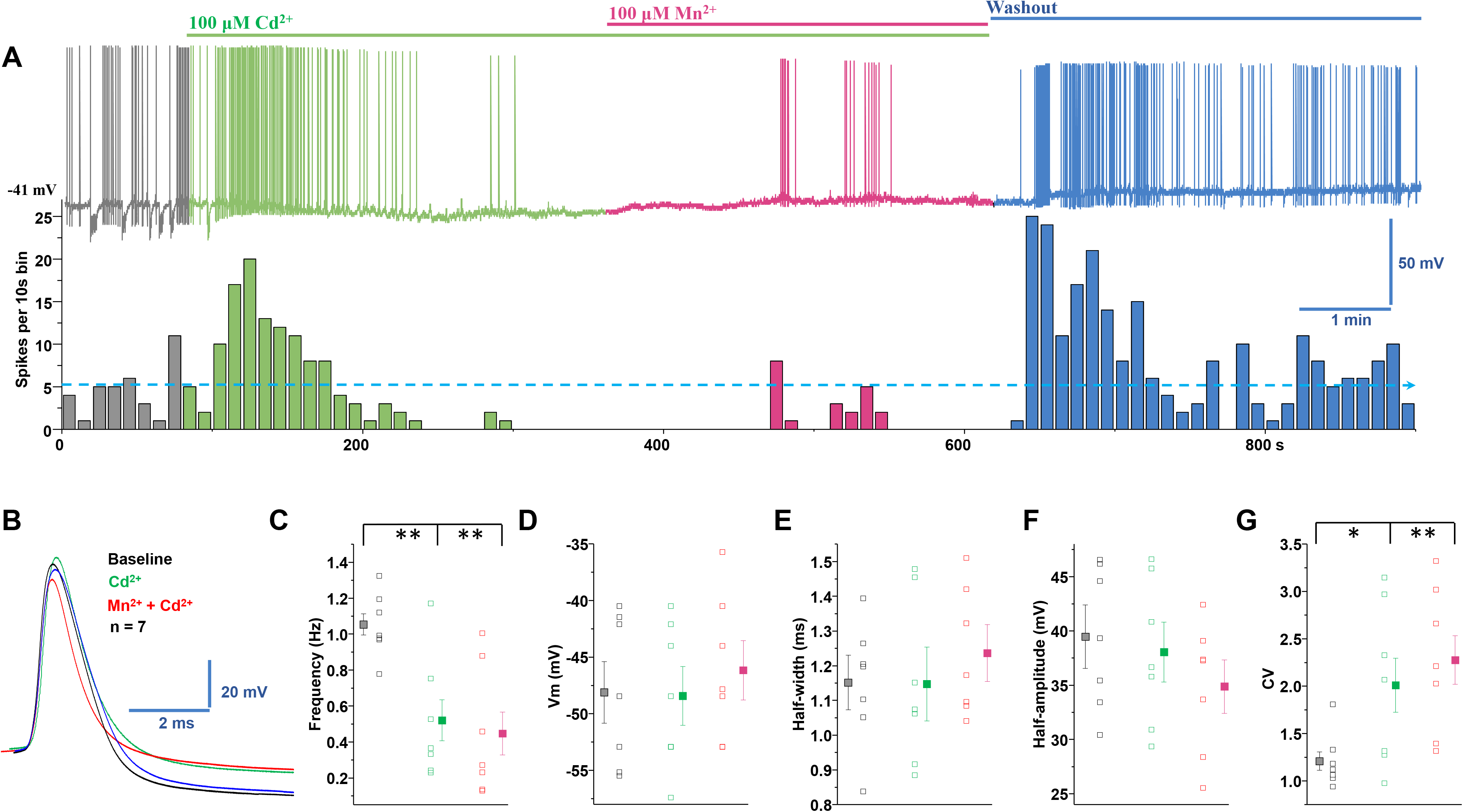
Cadmium blockade of voltage-gated Ca^2+^ channels inhibited manganese-stimulation of spontaneous firing activity of dopamine neurons. ***A,*** Top: representative spontaneous spike activity of a dopamine neuron before and after exposure to cadmium (Cd^2+^ a nonselective voltage-gated Ca^2+^ channel blocker) and Cd^2+^ + Mn^2+^. Cd^2+^-blockade of voltage-gated Ca^2+^ channels inhibited Mn^2+^-stimulation of spontaneous firing activity. Bottom: histogram of firing frequency from above trace showing firing frequencies were lowered in Cd^2+^ only condition and following or Cd^2+^ plus Mn^2+^, but return to baseline after washout. ***B**,* Superimposed representative single action potential traces shown in A: baseline (black trace), 100 μM Cd^2+^ (green trace), 100 μM Mn^2+^ + Cd^2+^ (red trace), and washout (blue trace). ***C**,* Spontaneous firing rate significantly decreased following application of Cd^2+^ or Cd^2+^ plus Mn^2+^ (*F*_(2,18)_ = 8.3, *p* = 0.002, one-way ANOVA followed by Tukey’s test, *n* = 7 / group). ***D**,* Application of Cd^2+^ or Cd^2+^ plus Mn^2+^ did not change the membrane potential. ***E**,* Half-width was measured at half-maximal voltage of action potential; there was no change in the half-width of, Cd^2+^ and Cd^2+^ plus Mn^2+^ experimental groups. ***F**,* Cd^2+^ or Cd^2+^ plus Mn^2+^ treatments did not change the amplitude of action potential. ***G**,* Both Cd^2+^ and Cd^2+^ plus Mn^2+^ treatments increased the coefficients of variation of the interspike interval (*F*_(2,18)_ = 4.6, *p* = 0.024, one-way ANOVA followed by Tukey’s test, *n* = 7 / group). *p < 0.05; **p < 0.01.

The Ca^2+^ current involved in pace-making activity is partly due to L-type currents, which are activated at subthreshold potential which might contribute to tonic firing (Putzier et al., 2009). To investigate the possible role of these channels, we asked whether nifedipine-blockade of L-type Ca^2+^ (Ca_v_1) channels inhibited manganese-stimulation of spontaneous firing activity of dopamine neurons. As shown in Figure 7, we found nifedipine inhibited manganese-stimulation of spontaneous firing activity. Consistent with the results in Figure 3-4, the firing frequencies were increased in the presence of manganese and Ca^2+^-free condition. Similar to the Cd^2+^-inhibition of manganese-stimulation of firing activity of dopamine neurons, co-application of nifedipine inhibited the manganese-stimulation of firing activity (Figure 7A, C: *F*_(2,18)_ = 44.4, *p* = 0.0006, one-way ANOVA followed by Tukey’s test, *n* = 7 group). Similar to Cd^2+^, the manganese in Ca^2+^– free solution condition or the co-administration of manganese plus nifedipine in Ca^2+^-free solution did not change the membrane potential (Figure 5D) or AP half-width (Figure 5B, E), but both manganese in Ca^2+^–free solution and additional application of nifedipine largely truncated AP amplitudes (Figure 7A, F: *F*_(2,18)_ = 11.80, *p* =0.001, one-way ANOVA followed by Tukey’s test, *n* = 7 group). The co-administration of manganese in Ca^2+^-free manganese and nifedipine significantly increased the CV (Figure 7G: *F*_(2,18)_ = 14.11, *p* = 0.0016, one-way ANOVA followed by Tukey’s test, *n* = 7 group). These unexpected results support the interpretation that manganese may influx into the neurons through the nifedipine-sensitive voltage-gated Ca^2+^ channels.

**Figure 7.**
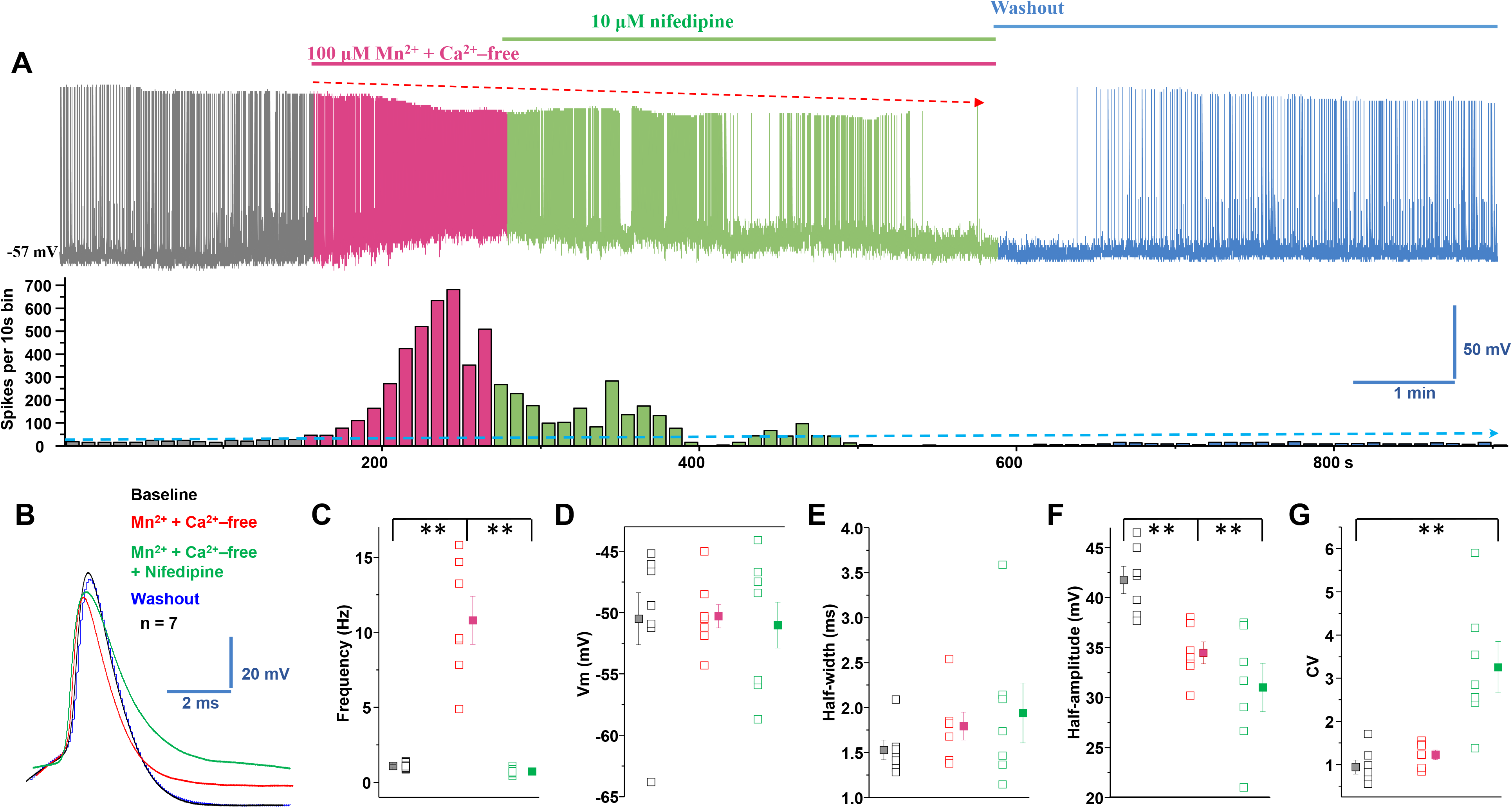
Blockade of L-type Ca^2+^ channels inhibited manganese-stimulation of spontaneous firing activity of dopamine neurons. ***A,*** Top: representative spontaneous spike activity of a dopamine neuron before and after exposure to Mn^2+^ plus Ca^2+^-free and nifedipine, an L-type Ca^2+^ channel blocker. Nifedipine-blockade of voltage-gated Ca^2+^ channels inhibited Mn^2+^-stimulation of spontaneous firing activity. Bottom: histogram of firing frequency from above trace showing firing frequencies were increased in combination of Mn^2+^ and Ca^2+^-free, lowered in additionally added nifedipine, but return to baseline after washout. ***B,*** Superimposed representative single action potential traces shown in A: baseline (black trace), Mn^2+^+ Ca^2+^-free (red trace), Mn^2+^+ Ca^2+^-free + nifedipine (green trace), and washout (blue trace). ***C,*** Mn increased firing rate even in the combination of Mn^2+^ and Ca^2+^-free. Additional application of nifedipine significantly reduced firing rate (*F*_(2,18)_ = 44.4, *p* = 0.0006, one-way ANOVA followed by Tukey’s test, *n* = 7 / group). ***D,*** Application of Mn^2+^+ Ca^2+^-free or further administrated nifedipine did not change the membrane potential. ***E,*** Half-width was measured at half-maximal voltage of action potential; there was no change in the half-width of, Mn^2+^+ Ca^2+^-free and additional plus nifedipine experimental groups. ***F,*** Both Mn^2+^ plus Ca^2+^-free and further added nifedipine condition significantly depressed the amplitude of action potential (*F*_(2,18)_ = 11.80, *p* = 0.001, one-way ANOVA followed by Tukey’s test, *n* = 7 / group). ***G,*** Mn^2+^ plus Ca^2+^-free and additional application of nifedipine increased the coefficients of variation of the interspike interval (*F*_(2,18)_ = 14.11, *p* = 0.0016, one-way ANOVA followed by Tukey’s test, *n* = 7 / group). **p < 0.01.

### Single neuron recording revealed manganese competes with Ca^2+^ influx and generates Ca^2+^ channel-like currents

Because the blockade of voltage-gated Ca^2+^ channels prevented the manganese-mediated increases in the spontaneous firing activity of dopamine neurons independent of the presence of intracellular or extracellular Ca^2+^, we then examined the hypothesis that voltage-gated Ca^2+^ channels on dopamine neurons where directly permeable to extracellular manganese. After replacing external Ca^2+^ with equimolar manganese, we observed manganese induced a Ca^2+^ channel-mediated current, which was evoked by a series of 200 ms depolarizing steps from −60 to +85 mV in 5 mV increments (Figure 8A_1_). The currents were composed of a rapidly inactivating transient component and a slowly inactivating persistent component. The current–voltage relationship of the manganese current is shown in Figure 8B. The manganese currents were activated at voltages around 20 mV and peaked at approximately 30 mV. Next, we asked whether the activity of Ca^2+^ channels in midbrain dopamine neurons is affected by these manganese-mediated currents. Since there is little information on whether manganese modulates Ca^2+^ currents in dopamine neurons, we focused on the action of manganese on whole-cell Ca^2+^ currents. Total Ca^2+^ currents (Figure 8A_2_, C) were activated at voltages around 20 mV and peaked at approximately 45 mV. Bath application of 100 μM manganese suppressed the peak amplitude of total Ca^2+^ currents (Figure 8A_3_, C, *\*p* < 0.05, two-tailed Student’s *t* tests; *n* = 7 / group). These data suggest in dopamine neurons, manganese might compete with Ca^2+^ influx and generate Ca^2+^ channel-gated-like currents that in turn alter the activity of the neurons. To our knowledge, these are the first reported findings that Ca^2+^ channels may be directly permeable to manganese and may represent a distinct role of manganese in regulating neuronal activity.

**Figure 8.**
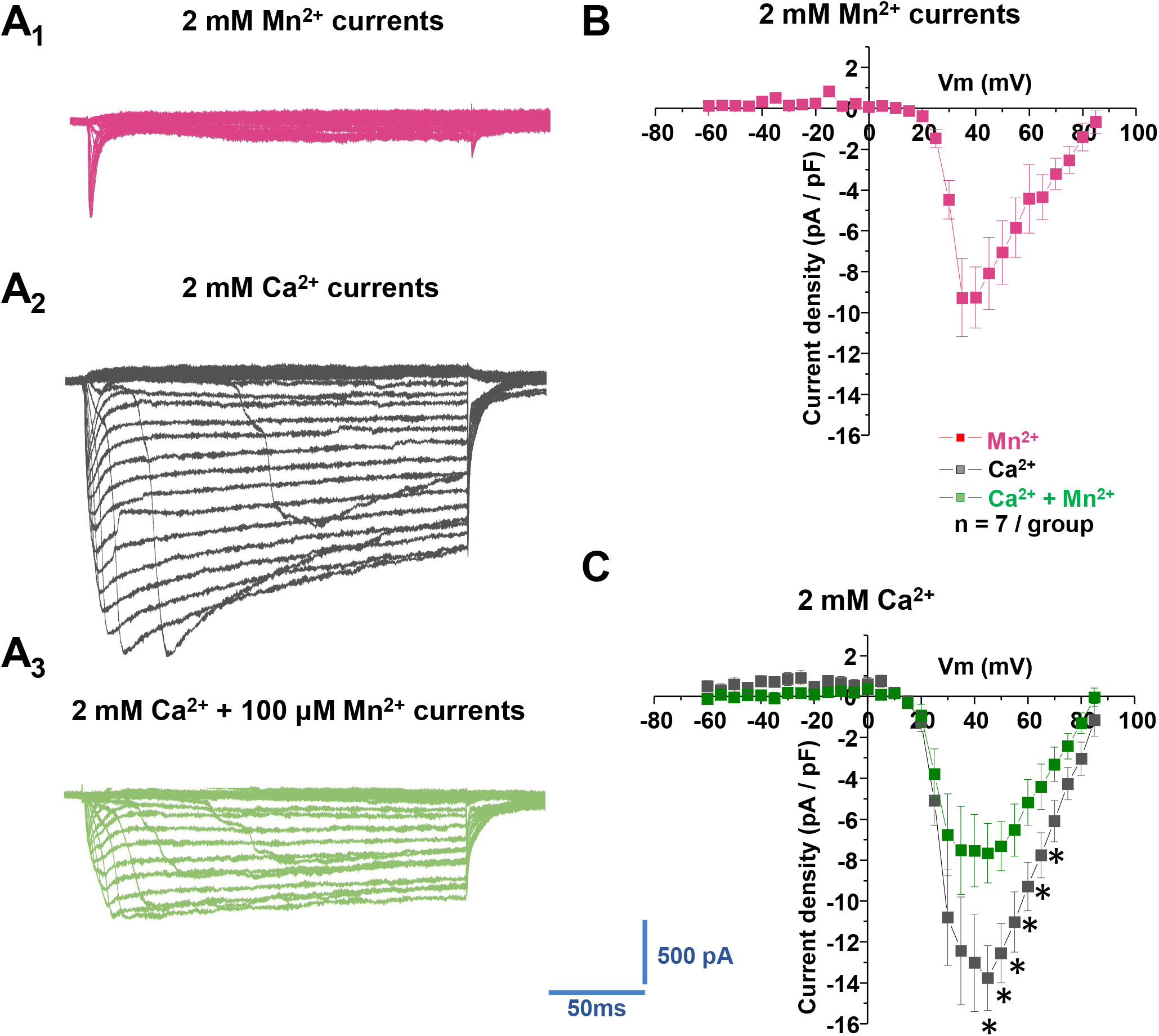
Manganese reduced voltage-gated Ca^2+^ currents. ***A,*** Representative traces of inward currents in response to voltage steps (5 mV, 250 ms) from −60 mV to +85 mV in a bath solution containing 0.5 μM TTX, 35 mM TEA and 1mM CsCl. ***A_1_***, Inward currents evoked by voltage steps during application of extracellular Ca^2+^-free plus 2mM Mn^2+^ solution. ***A_2_***, Inward currents evoked by voltage steps in the presence of 2 mM Ca^2+^. ***A_3_***, Inward currents evoked by voltage steps in the presence of 2 mM Ca^2+^ plus 100 mM Mn^2+^. ***B,*** Plot shows the current-voltage relationship (I–V curves) induced by Mn^2+^. ***C,*** Plot shows current-voltage relationships (I–V curves) of peak voltage-gated Ca^2+^ current density in dopamine neurons at baseline (black squares) and following Mn^2+^ exposure (blue squares), (*\*p* < 0.05, two-tailed Student’s *t* tests; *n* = 7 / group).

### Manganese-enhanced magnetic resonance imaging provides evidence for an Ca^2+^-channel dependent mechanism in the midbrain *in vivo*

To further examine the manganese interaction with Ca^2+^ channels, we asked whether manganese can influx into the Ca_v_1.2 and Ca_v_1.3 expressing cells and whether nifedipine inhibits manganese influx through Ca_v_1 channels. To examine this hypothesis *in vivo*, we performed manganese enhanced magnetic resonance imaging (MEMRI) in animals pretreated with saline or manganese. I.p. injections of 70 mg/kg manganese has previously been shown to significantly increase manganese levels in the basal ganglia (Dodd et al., 2005), and also is a side-effect of contrast agent for MRI (Poole et al., 2017). Manganese frequently serves as a neuronal contrast agent to enhance functional brain mapping at a higher spatial resolution than typical fMRI studies (Lee et al., 2005). In this study, we utilized MEMRI and T_1_ mapping to study basal levels of brain activity before and after nifedipine-blockade of voltage-gated Ca^2+^ channels (Figure 9 and Supplemental Figure 3). A commonly used concentration of nifedipine (15 mg/kg) (Morellini et al., 2017, Giordano et al., 2010) was injected before 30 min of manganese exposure. As expected, we found that T_1_ relaxation (quantified as R_1_ or the rate of T_1_ relaxivity in s^-1^) following manganese treatment exhibited predominantly higher (faster) R_1_ in ventral tegmental area (VTA) and substantia nigra (SN) compared to other cortical and subcortical regions (Supplemental Figure 3, *F*_(8,72)_ = 2.35, *p* = 0.026, one-way ANOVA followed by Tukey’s test, test, *n* = 8 – 9 / group). This reflects a shortening of VTA and SN T1 relaxation time due to the accumulation of the paramagnetic manganese. Importantly, and consistent with single neuron recordings (Figure 6, 7, and 8), we observed that nifedipine-blockade of L-type Ca^2+^ channels reduced T_1_ shortening effects of manganese in dopaminergic midbrain regions (Fig 9, VTA: *t_(15)_* = −2.2, p = 0.04, two-tailed Student’s t tests; Sn: *t_(15)_* = −2.7, p = 0.015, two-tailed Student’s t tests; *n* = 8 – 9 / group). This was observed as a slower rate R_1_ in the nifedipine/manganese group relative to manganese only. Therefore, both *in vitro* (single neurons recording) and *in vivo* studies suggest that nifedipine decreases manganese-stimulation of neuronal activity and its accumulation in the multiple brain regions including the dopamine neuron enriched midbrain.

**Figure 9.**
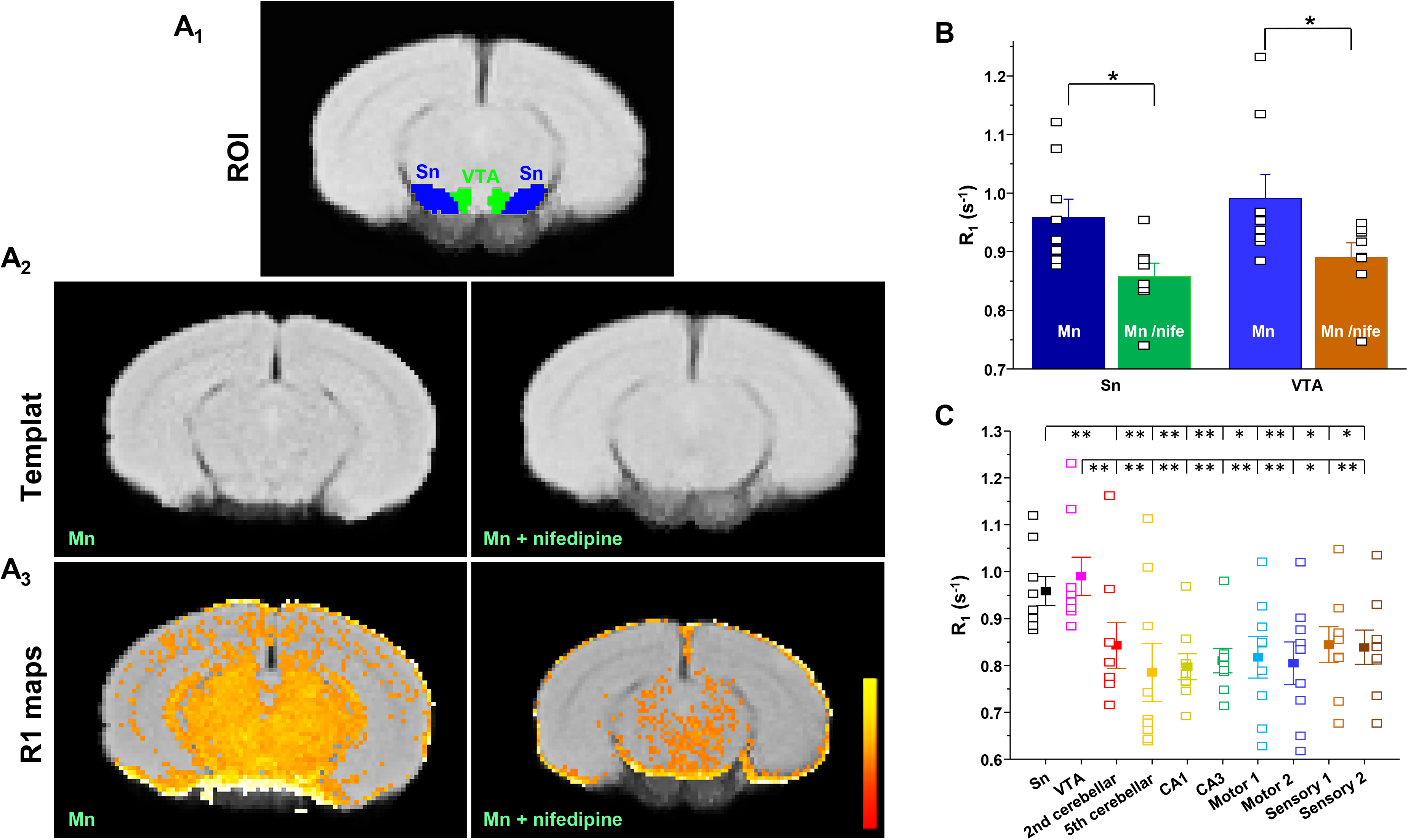
Manganese enhanced magnetic resonance imaging (MEMRI) provides evidence of an *in vivo* Ca^2+^-channel dependent mechanism involved in midbrain manganese accumulation. R_1_ (1/T_1_) maps. The top figure displays the location of the ventral tegmental area (VTA) and substantia nigra (SN). The middle figure displays a custom template from all controls and experimental groups (nifedipine administered rats). The bottom figure displays the average R_1_ map of both cohorts. Maps display values up to 2 s^-1^ with yellow representing large values (∼2 s^-1) and red representing low values (∼ 0). The control group shows a small number of voxels of R_1_ larger within the threshold showing less Mn^2+^ infiltration in the midbrain. *A_1_*, Mouse brain atlas-based segmentation of the VTA and SN. ***A_2_***, Template brain for Mn alone or with nifedipine treatment that were aligned with mouse brain atlas. ***A_3_***, Parametric maps of T_1_ relaxation rate (R_1_ in msec^-1^) show that calcium channel blockade with nifedipine treatment reduces R_1_ in midbrain and surrounding areas. Scale bar indicates intensity of R_1_. ***B***, Nifedipine reduces T_1_ relaxation rate (R_1_) in VTA and SN (VTA: *t_(15)_* = −2.2, p = 0.04, two-tailed Student’s t tests; Sn: *t_(15)_* = −2.7, p = 0.015, two-tailed Student’s t tests; *n* = 8 – 9 / group). ***C***, A greater Mn^2+^ accumulation produces faster rates of T_1_ relaxation (R_1_) in VTA and SN than in other cortical and subcortical nuclei (*F*_(9,72)_ = 3.96, *p* < 0.001, one-way ANOVA followed by Tukey’s test, *n* = 8 – 9 / group). **p < 0.01.

### Manganese influxes into Ca_v_1.2 and Ca_v_1.3 expressing cells in a nifedipine-dependent manner

Our data so far supports the interpretation that manganese may influx via nifedipine-sensitive Ca^2+^ channels. To examine this idea directly in a reduced system, HEK cells were transiently transfected with either a control empty vector or Ca_v_1.2 or Ca_v_1.3 cDNA. Western blot analysis confirmed the greater expression of Ca_v_1.2 or Ca_v_1.3 in the transfected cells (Figure 10A). Overexpression of Ca_v_1.2 increased the uptake of ^54^Mn at 5 and 15 min after the addition of ^54^Mn, while there was no difference observed between the Ca_v_1.3 expressing and control cells (Figure 10B). Uptake of ^54^Mn by HEK cells overexpressing Ca_v_1.2 was inhibited by pretreating with nifedipine (Figure 10C). These findings support the interpretation that manganese regulates dopamine neuronal activity and manganese influx via a nifedipine-sensitive Ca_v_1.2-channel-mediated mechanism.

**Figure 10.**
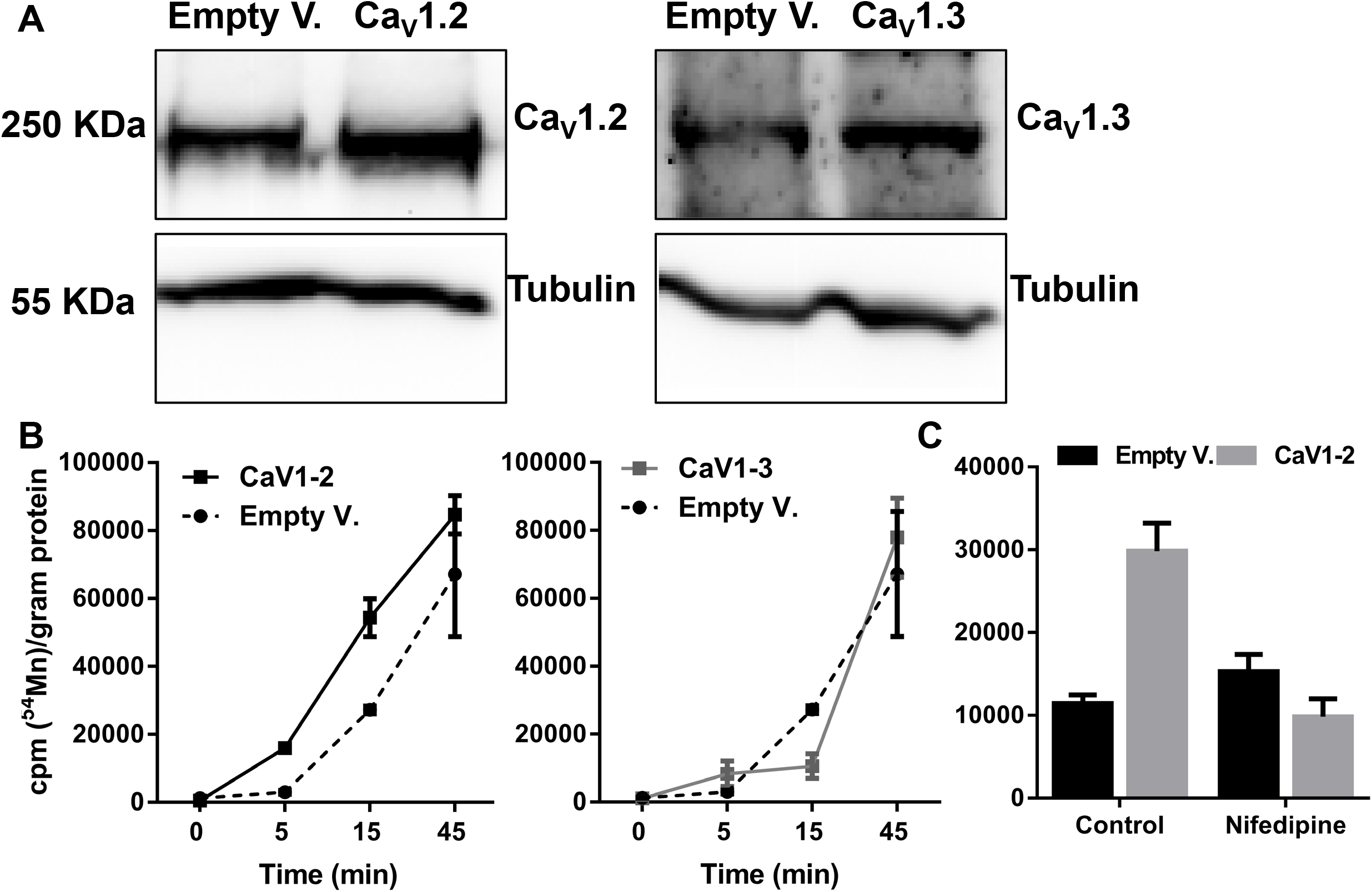
^54^Mn uptake is greater in Ca_v_1.2 overexpressing HEK293 cells. ***A,*** Representative western blot analysis is showing the expression of Ca_v_1.2 and Ca_v_1.3 in cells transiently-transfected to overexpress Ca_v_1.2 and Ca_v_1.3. ***B,*** Greater ^54^Mn uptake is shown in Ca_v_1.2 overexpressing cells at the indicated time points. ***C,*** Nifepidine inhibits ^54^Mn uptake at 15 minutes in Ca_v_1.2 overexpressing cells. (One-way ANOVA followed by Tukey’s test, n = 3 / group). *p<0.05)

### In silico modeling support the possibility that manganese can interact with the Ca^2+^ binding site in the Ca_v_1 channel

Because the crystal structure for Ca_v_1.2 is not available, Cav1.1 was used for the homology modeling studies to investigate whether the putative structure of Ca_v_1.2 could informs us about the manganese cation transport. First, the available electron microscopy structure for the homologous Ca_v_1.1 channel was used to build a homology model using the SWISS-MODEL program (Figure 11) (Waterhouse et al., 2018). The helical regions of the mouse Ca_v_1.1 structure (Wu et al., 2016) and the helical regions for the Ca_v_1.2 model were compared by protein BLAST (Altschul et al., 1990) and found to be 72% identical and 81% similar. Further, when all amino acids within 7 Å of the bound Ca^2+^ atom found in the Ca_v_1.1 structure were selected and compared to amino acids in the model structure for Ca_v_1.2, we found 100% identity in this region, including the carboxylate ligands for the Ca^2+^ atom. We conclude the Ca_v_1.2 channel has an extremely similar structure and presumably cation-binding capability when compared with Ca_v_1.1. Because the ionic radius of Mn^2+^ is smaller than for Ca^2+^ (Persson, 2010), it should be adequately accommodated by the channel. Taken together, the structural features of the Ca_v_1.2 channel and our experimental findings argue for the possibility of the Ca_v_1.2 channel conducting a manganese ion. While these data collectively suggest manganese produces a Ca^2+^ channel-mediated current in dopamine neurons, which increases rhythmic firing activity of dopamine neurons, the connection between manganese-mediated current and increased firing activity of dopamine neurons remain unclear. The next set of experiments are designed to determine the mechanism.

**Figure 11.**
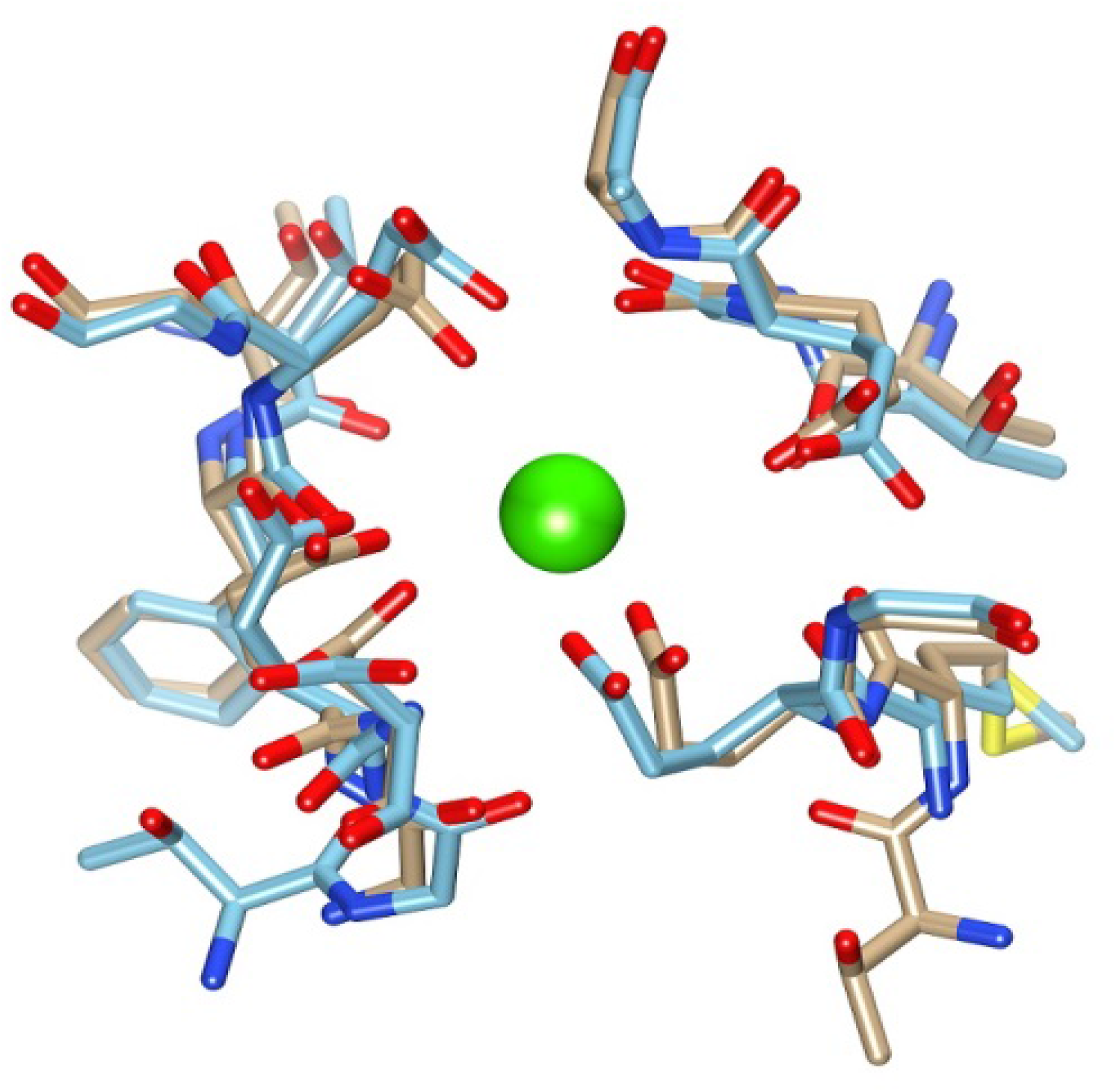
Superposition of msCa_v_1.1 Ca binding site structure with the msCa_v_1.2 model. The Ca^2+^ ion (green) as located in the msCa_v_1.1 structure (tan color backbone) (PDB ID 5GJW). The homology model structure for Ca_v_1.2 alpha chain is shown in the blue color backbone.

### Blockade of large-conductance Ca^2+^-activated potassium (BK) channels decreased the spontaneous firing activity and prevented manganese-stimulation of firing frequency

Previously, we have shown that paxilline-blockade of BK channels reduces the spontaneous firing activity, broadens the width and enhances the amplitude of action potentials in midbrain dopamine neurons (Lin et al., 2016). Since Mn increased the firing frequency, decrease the width and the amplitude of action potentials (Figure 1), we postulated that manganese-stimulation of the firing frequency, narrowing of action potential width and reduction of action potential amplitude might be due to BK channel activation. Consistent with our previous reports, paxilline suppressed the firing frequency of midbrain dopamine neurons (Figure 12A, B). Pre-treatment with paxilline abolished the manganese-induced increase in firing frequency (Figure 12A, C: baseline: 1.5 ± 0.2 Hz vs. Mn + paxilline: 0.51 ± 0.1 Hz; *F*_(2,20)_ = 11.2, *p* = 0.0005, one-way ANOVA followed by Tukey’s test; *n* = 8 / group) and inhibited manganese’s modulation of action potential amplitude (Figure 12A, F). Comparison of the firing frequency after treatment with paxilline alone or paxilline + manganese treatment revealed a further reduction in the firing frequency (Figure 12C; paxilline: 0.81 ± 0.1 Hz). Consistent with our previous report (Lin et a., 2016), blockade of BK channels depolarized membrane potential (Figure 12D; Baseline: −45.1 ± 1.6 mV vs paxilline: - 40.1 ± 1.8 mV; *F*_(2,21)_ = 3.5, *p* = 0.04, one-way ANOVA followed by Tukey’s test; *n* = 8 / group), broadened AP half-width (Figure 12B, E: baseline: 1.3 ± 0.1 ms vs. paxilline: 1.8 ± 0.2 ms;; *F*_(2,21)_ = 5.9, *p* = 0.009, one-way ANOVA followed by Tukey’s test; *n* = 8 / group), increased action potential amplitude (Figure 12B, F; baseline: 28.1 ± 1.6 mV vs. paxilline: 32.6 ± 1.1 mV; *F*_(2,21)_ = 6.6, *p* = 0.005, one-way ANOVA followed by Tukey’s test; *n* = 8 / group) and increased the CV (Figure 12G; baseline: 1.8 ± 0.2 vs. paxilline: 2.8 ± 0.3; *F*_(2,21)_ = 2.8, *p* = 0.015, one-way ANOVA followed by Tukey’s test; *n* = 8 / group). After pretreatment with paxilline, we found manganese did not reverse paxilline-induced membrane depolarization (Figure 12D), but it did diminish the effects of paxilline on the action potential half-width (Figure 12E), amplitude (Figure 12F) and CV (Figure 12G). These results suggest that the manganese-stimulation of spontaneous spike activity of dopamine neurons is possibly due to BK channels activation.

**Figure 12.**
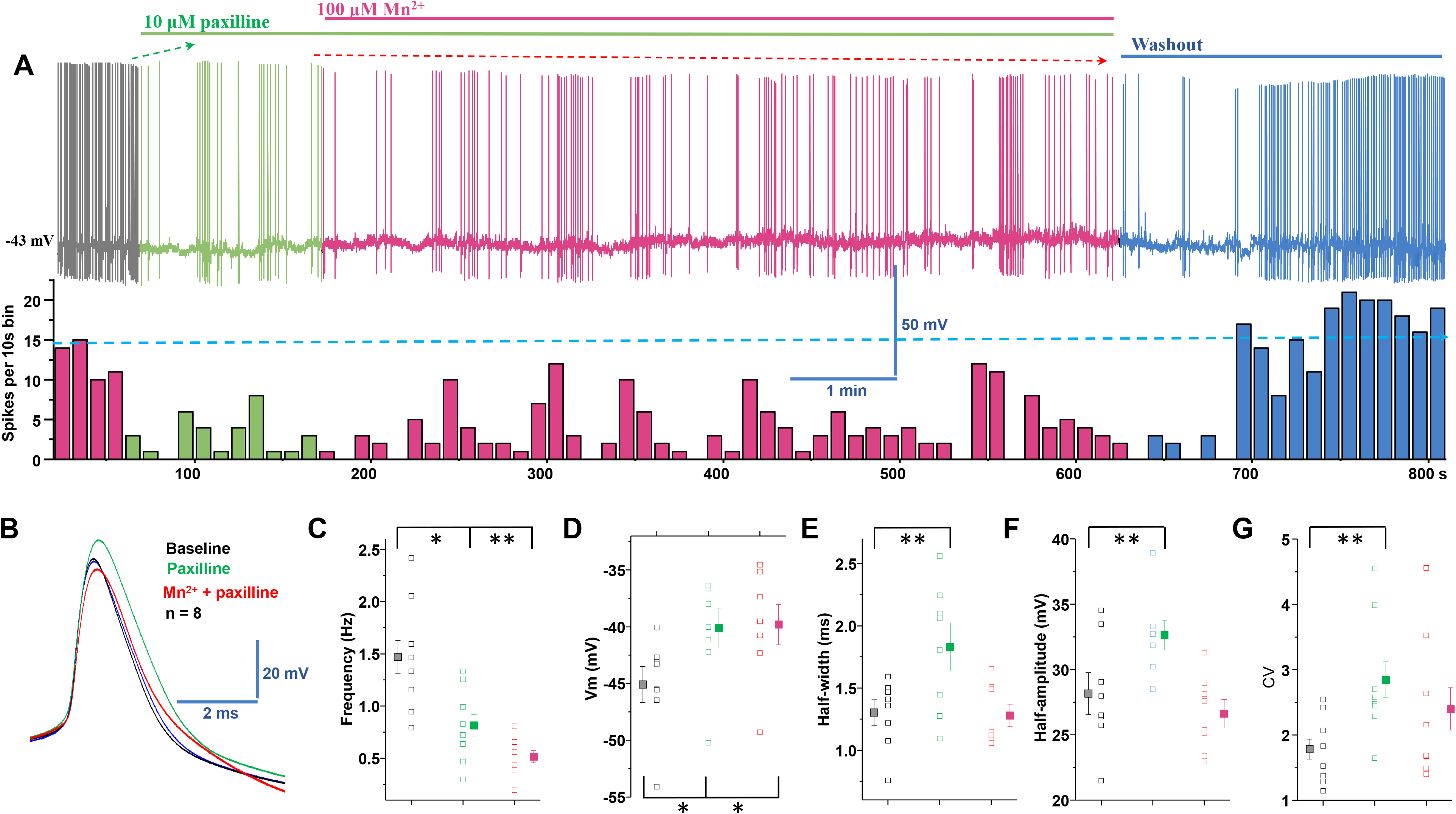
Blockade of BK channels inhibited Manganese-stimulation of the spontaneous firing activity in dopamine neurons. ***A,*** Top: representative spontaneous firing activities of a dopamine neuron exposed to paxilline followed by Mn treatment. Firing rate returns to baseline after washout with artificial cerebral spinal fluid (aCSF) solution. Bottom: Rate histogram of above trace. ***B,*** Superimposed traces of representative single action potential shown in A: baseline (black trace), paxilline (green trace), Mn (red trace) and washout (blue trace). ***C,*** Paxilline reduced the spontaneous firing rate; Mn treatment did not attenuate the paxilline-induced reduction of firing activity. ***D,*** Paxilline and concomitant Mn + paxilline applications significantly depolarized the membrane potential. ***E,*** Half-width is measured at half-maximal voltage of action potential. The half-width was broadened after paxilline application, but returned to baseline after co-administration of Mn and paxillin. ***F,*** Paxilline increased the amplitude of action potential, but diminished the effect of Mn on the action potential amplitude. ***G,*** Blockade of BK channels exhibited larger coefficients of variation of the interspike interval. *p < 0.05; **p < 0.01, n = 8 per group.

### Manganese increases membrane expression of BK-α subunits

The BK-α subunit is the pore-forming unit of the BK channel (Knaus et al., 1994). There is a direct relationship between BK channel regulation of neuronal excitability and the level of BK-α subunits at the surface membrane (Shruti et al., 2012; Chen et al., 2013; Cox et al., 2014). Since blockade of BK channels abolished the effect of manganese-stimulation of the spontaneous firing frequency of dopamine neurons, we examined the possibility that manganese-mediated enhancement of neuronal excitability is due to increased BK-α subunit levels at the plasma membrane. To test this, we used TIRF microscopy to monitor fluorescently-tagged BK channel trafficking. The TIRF profile of GFP-α subunits was examined in HEK293 cells expressing GFP-α subunits in the presence or absence of 100 μM manganese. As shown in Figure 13 and supplemental movie 4, the GFP-α subunits’ fluorescence signal at the membrane remained stable in the vehicle (external solution)-treated control group. The TIRF profile of GFP-α subunits was markedly increased following manganese treatment (*F*_(1,164)_ = 163.8, †: *p* < 0.01, one-way ANOVA followed by Tukey’s test, *n*: baseline = 7; Mn = 8). Because TIRF microscopy detects fluorophores at or near the plasma membrane, these results indicate that manganese’s stimulation of neuronal excitability is, in part, due to manganese-induced increases in surface BK channel expression and thus activity. To test this possibility, we used excised inside-out patches to record currents in response to test voltage steps. Figure 14 shows representative BK currents in response to test voltage steps in the absence or presence of manganese. The I–V curves of these recordings demonstrate Mn significantly increased peak amplitude of BK currents (*\*p* < 0.05, two-tailed Student’s *t* tests, *n* = 6 / group). Taken together, these results suggest manganese enhances BK channel activity that we have previously shown increases firing activity of dopamine neurons (Lin 2016).

**Figure 13.**
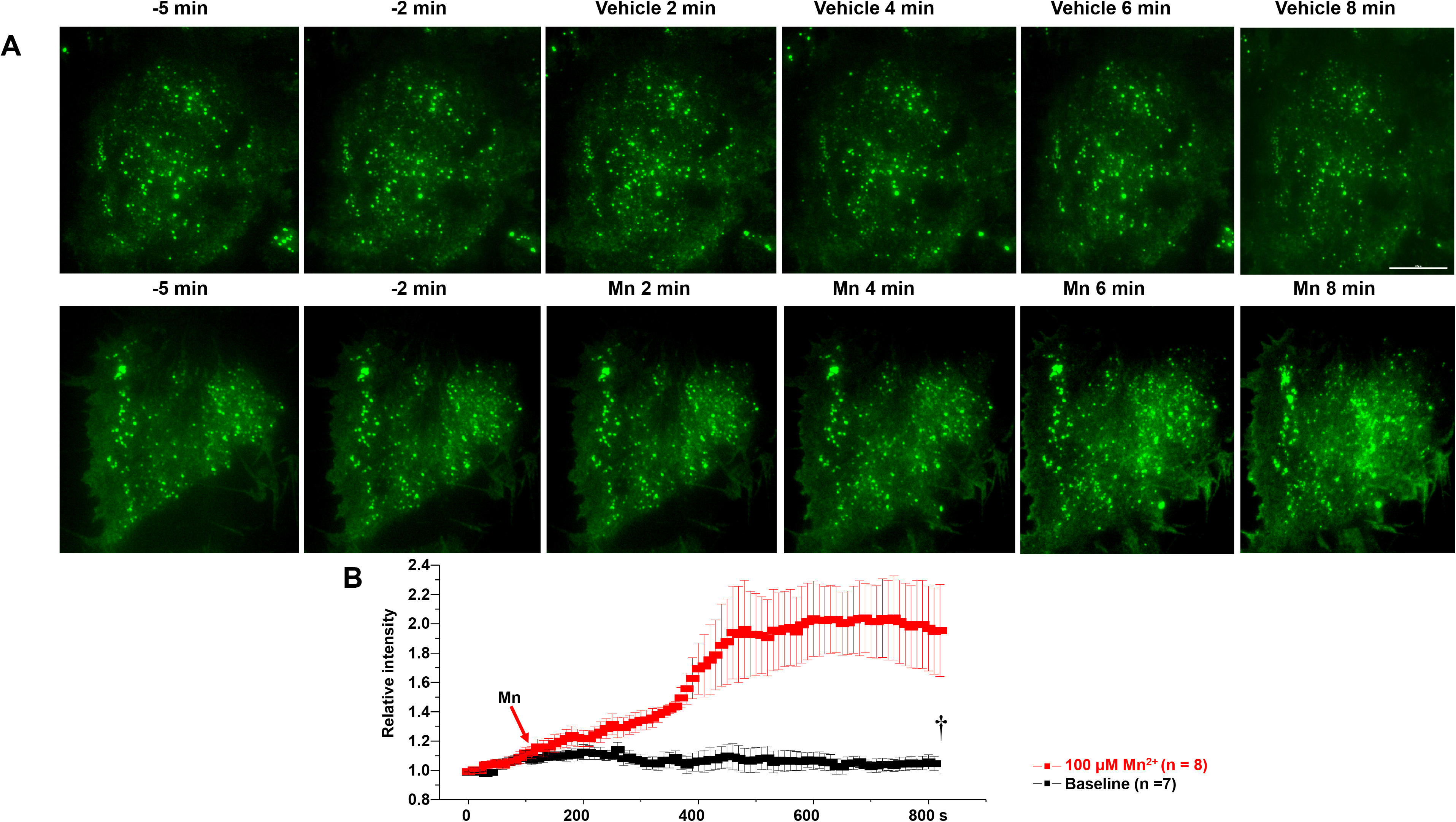
Membrane localization of BK GFP-α subunit is increased followed manganese exposure. ***A,*** Representative TIRF microscopy images of GFP-α subunit following vehicle or 100 μM Mn^2+^ treatments. ***B,*** Analyses of relative fluorescence intensities at the surface membrane following vehicle or 100 μM Mn^2+^. Scale bar: 20 μm. †: p < 0.01.

**Figure 14.**
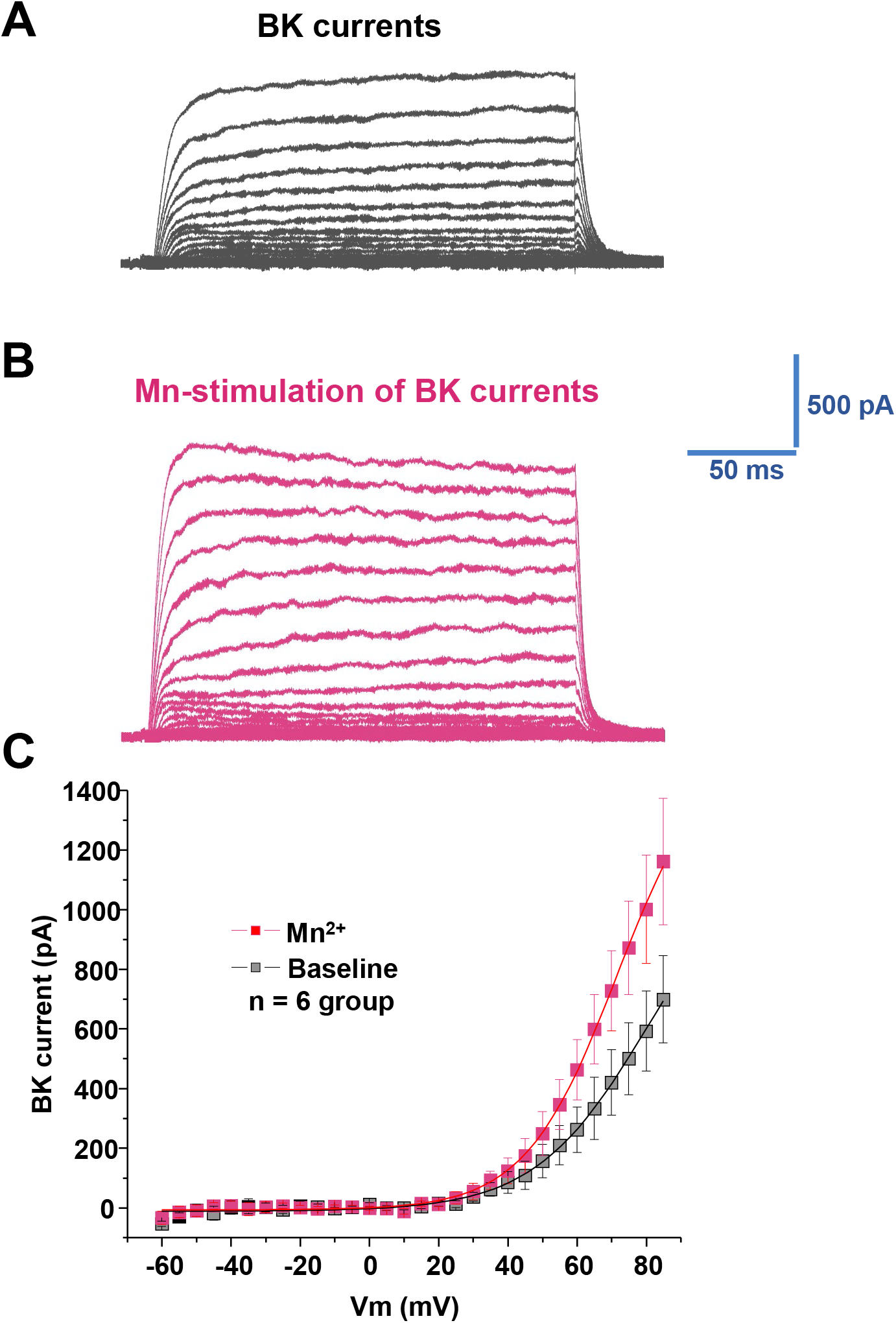
Manganese increases BK currents. ***A,*** Representative traces of outward currents in response to voltage steps (5 mV, 250 ms) from −60 mV to +85 mV in a bath solution containing 0.5 μM TTX, 35 mM TEA and 1mM CsCl. ***A***, In cells expressing BK-α subunits, families of outward currents were evoked by voltage steps from −60 to +85 mV for 250 ms with 5 mV increments every 5 s from the holding potential of −70 mV, before (A) and after (B) Mn administration. ***C,*** shows peak current-voltage relationships (I–V curves) before (baseline) and after Mn application. *p < 0.05.

## Discussion

Prolonged exposure to low levels of manganese or single large exposure results in its accumulation in multiple brain regions leading to dysfunction of central nervous system and an extrapyramidal motor disorder referred to as manganism (Aschner et al., 2009). In this study, we used multiple complementary approaches to examine the manganese-regulation of dopaminergic excitability and to determine the potential mechanisms involved. We found manganese increased the spontaneous firing activity of dopamine neurons, decreased the amplitude and half-width of action potentials, and reduced the variation of interspike interval. Unexpectedly, neither the removal of extracellular Ca^2+^ and/or the chelation of intracellular Ca^2+^ modulated the manganese-stimulation of spontaneous firing frequency of dopamine neurons. Live cell two-photon imaging of GCaMP6f expressing dopamine neurons support the electrophysiology data showing no change in intracellular Ca^2+^ levels after manganese application. In contrast, we found manganese-regulation of dopamine neurons was blocked by cadmium, a nonselective Ca^2+^ channel blocker or by nifedipine, an L-type Ca^2+^ channel blocker, suggesting manganese can potentially influx through voltage-gated Ca^2+^ channels. Furthermore, we identified a Ca^2+^ channel-mediated manganese current that reduced voltage-gated Ca^2+^ currents, supporting the idea that manganese may compete with Ca^2+^ influx, leading to activation of BK channels and increased spontaneous firing activity of dopamine neurons.

### Manganese competes with Ca^2+^ entry through voltage-gated Ca^2+^ channels to enhance excitability of dopamine neurons

The action potential of dopamine neurons is slow and broad, which maximizes Ca^2+^ entry and promotes slow rhythmic activity (Bean, 2007). The slow, rhythmic activity (2–10 Hz) in these neurons is autonomously generated and accompanied by slow oscillations in intracellular Ca^2+^ concentration that are triggered by the opening of plasma membrane Ca_v_1 (Ca_v_1.2, Ca_v_1.3) Ca^2+^ channels and release of Ca^2+^ from intracellular, endoplasmic reticulum stores (Guzman et al., 2009; Morikawa and Paladini, 2011; Nedergaard et al., 1993; Puopolo et al., 2007). Thus, we tested the hypothesis that manganese increases Ca^2+^ entry into the neuron and/or releases Ca^2+^ from intracellular Ca^2+^ stores. Unexpectedly, our data support neither of these possibilities. We found performing the experiments in either extracellular Ca^2+^-free condition or chelation of intracellular Ca^2+^ did not impair manganese-stimulation of firing frequency or reduction of action potential amplitude. Consistently, live cell two-photon Ca^2+^ imaging showed manganese did not alter intracellular Ca^2+^ concentration in the midbrain dopamine neurons. Thus, although manganese has been previously shown to regulate Ca^2+^ homeostasis in astrocytes (Xu et al., 2009), this effect is not present in dopamine neurons. Previous studies suggest manganese can enter cells through a number of transporters (Gunshin, 1997; Lockman et al., 2001; Crossgrove and Yokel, 2005; Goytain et al., 2008; Itoh et al., 2008; Gunter et al., 2013). In addition, as a divalent cation, manganese may potentially target Ca^2+^ channels, which are also permeant to other divalent cations such as barium (Bourinet et al., 1996). Therefore, if manganese competes with Ca^2+^ entry at the level of voltage-gated Ca^2+^ channels, then a nonselective Ca^2+^ channel blocker such as cadmium (Cd^2+^) or a L-type Ca^2+^ channel nifedipine should block the effect of manganese on the firing frequency of dopamine neurons and action potential morphology. As shown in Figure 6 and 7, both of Cd^2+^ and nifedipine suppressed manganese–stimulation of spontaneous firing activities, which is consistent with recent reports showing dihydropyridine L-type channel inhibitors slow pacemaker activity of dopamine neurons at submicromolar concentrations (Nedergaard et al., 1993; Puopolo et al., 2007).

### Blockade of L-type Ca^2+^ (Cav1) channels decreases manganese influx

Since the 1980s, manganese has been used as a tool to increase the signal to noise ratio in magnetic resonance imaging (MRI) (Lauterbur et al., 1980). Consistent with previous reports (Aoki et al., 2004; Lee et al., 2005), following systemic manganese administration, manganese is accumulated in multiple brain regions including dopaminergic neuron enriched brain regions such as substantia nigra and ventral tegmental area (Figure 9 and Supplemental Figure 3). Here we found, nifedipine-blockade of L-type Ca^2+^ channels decreased manganese accumulation in several brain regions as well as manganese uptake in the Ca_v_1.2 expressing HEK cells. These data support the interpretation that manganese might be permeable through the Ca^2+^ channels. Consistent with this idea, the chemical properties of aqueous Ca^2+^ and Mn^2+^ ions and the smaller ionic radius of Mn^2+^ than for Ca^2+^ also support the hypothesis that manganese might be accommodated by the Ca_v_1 channels. While the crystal structure of Ca_v_1.2 is yet to be determined, *in silico* analyses of the homology model of the helical regions of the mouse Ca_v_1.1 structure and the helical regions for the Ca_v_1.2 were found to be 72% identical and 81% similar for Ca_v_1.2; therefore, it is possible that Mn^2+^ enters Cav expressing cells. Whether manganese is permeable to other Ca^2+^ channels in dopamine or other neuronal types requires further examination.

Manganese influx through L-type Ca^2+^ channels was further supported by our manganese enhanced magnetic resonance imaging (MEMRI) data (Figure 9). It should be noted that both neuronal and glial cells express Ca_v_1 channels and therefore the MEMRI and the nifedipine-inhibition of this response reported in our studies do not distinguish the cell type. While the effect of manganese on Ca_v_1.2 is consistent across other CNS cell types requires further investigation, our observations of manganese accumulation in the dopamine enriched brain region and *in vitro* studies on the effects of manganese on dopaminergic neurons could establish a biological basis for movement disorders associated with manganism, which typically include symptoms related to dysregulation of the dopaminergic system. Furthermore, although the MEMRI data suggest acute manganese exposure can be detected in multiple brain regions, dopaminergic brains regions, specifically the VTA and SN, displayed a greater enhancement of MRI signal by manganese compared to other brain regions observed in this study. Overall, we found that blockade of Ca_v_1.2 can decrease manganese regulation of dopamine neuron activity and its accumulation in the brain.

### Manganese increases membrane expression of BK channels α-subunit leading to enhanced BK channels activity

Structurally, BK channel complexes contain the pore-forming α subunit (four α subunits form the channel pore) and the regulatory β subunits (Knaus et al., 1994; Brenner et al., 2000; Lu et al., 2006). The intrinsic gating properties of BK channels are dynamically modulated by various kinases (White et al., 1991, Weiger et al., 2002) that regulate BK channels trafficking to the membrane (Chae and Dryer, 2005, Toro et al., 2006). Recently, we have shown that PKC activation decreases membrane expression of GFP-α subunits of BK channels (Lin et al., 2016). These data combined with reports showing manganese-activation of protein kinase delta triggers apoptosis in dopaminergic neurons (Kitazawa et al., 2005), led us to ask whether Mn increases GFP-α subunit membrane expression. We found manganese enhanced surface trafficking of BK channels (Fig 13) and increased BK channels activity (Fig 14). These data collectively describe a cellular mechanism for the paxilline-blockade of Mn-stimulated increases in firing frequency as well as its effect on the action potential morphology. Future studies will determine the PKC subtype/s involved in Mn-induced trafficking of GFP-α subunit of BK channels and the potential involvement of other BK channels subunits such as regulatory β subunits. Collectively, these findings reveal the cellular mechanism for manganese-regulation of dopamine neurons and reveal a unique therapeutic target to attenuate the untoward consequence of manganese exposure.

## Acknowledgements

This study was supported by DA026947, NS071122, OD020026, DA026947S1, DA043895

We are very grateful to Dr. Gregory Hockerman for providing plasmids for Ca_v_1.2-GFP and Ca_v_1.3-GFP, Dr. Richard B. Silverman for providing HEK 293 cells stably expressing Cav1.2 or 1.3 and Dr. Robert Brenner for providing HEK293 cells stably expressing GFP-BK-α subunits.

## Figure Legends

**Supplemental Figure. 1.**
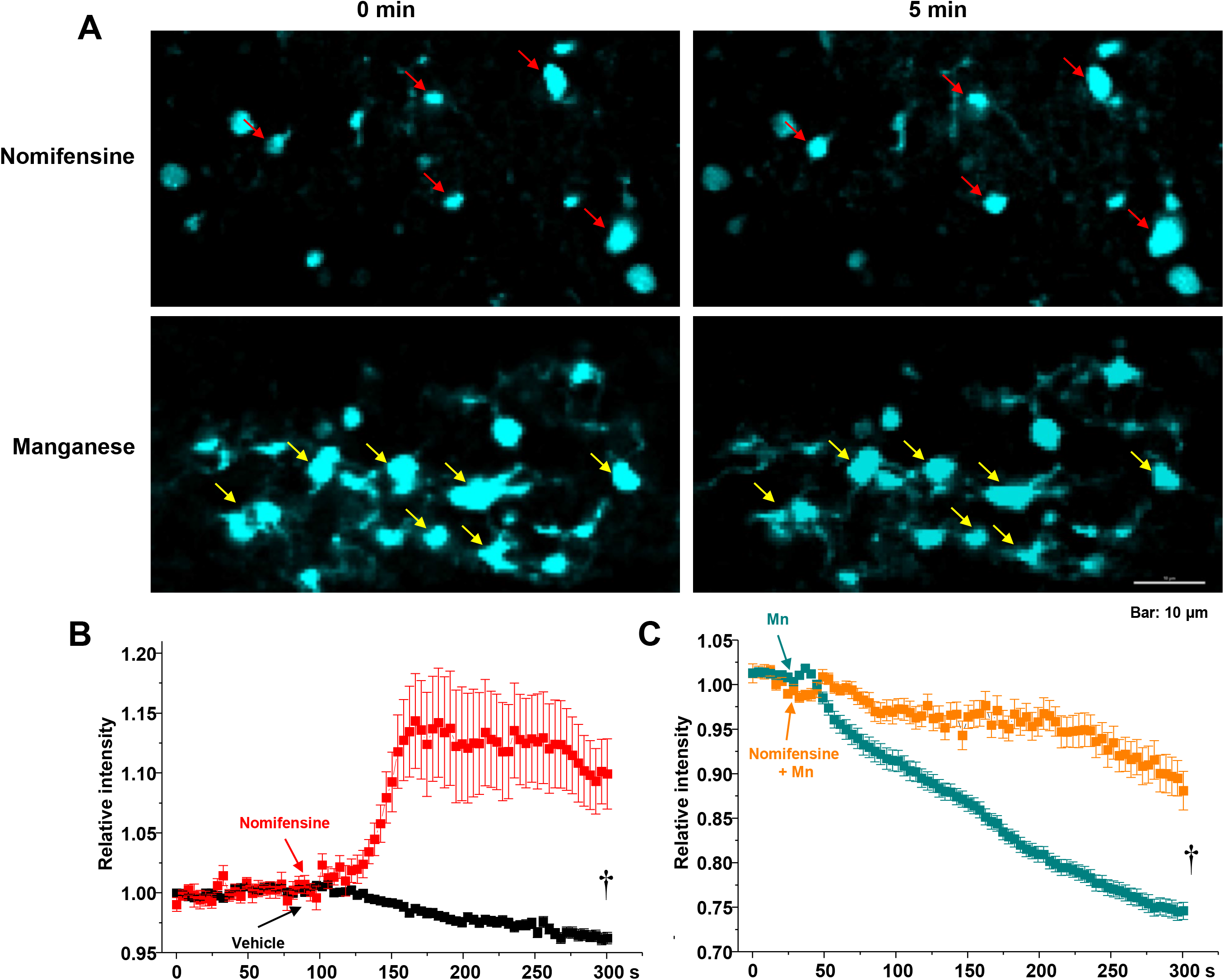
Mn^2+^ induces release of FFN200 from dorsal striatum. ***A***. Time lapse images of a striatal slice loaded with FFN200 following nomifensine (A top panel) or 100 μM Mn^2+^ treatments (A bottom panel). Mn^2+^ treatment decreased FFN200 puncta intensities (yellow arrows). ***B***. ***C***. Quantification of FFN200 puncta fluorescence normalized to the averaged intensity of first 60sec for each experimental condition. Scale bar: 10 μm. †: p < 0.01.

**Supplemental Figure 2.**
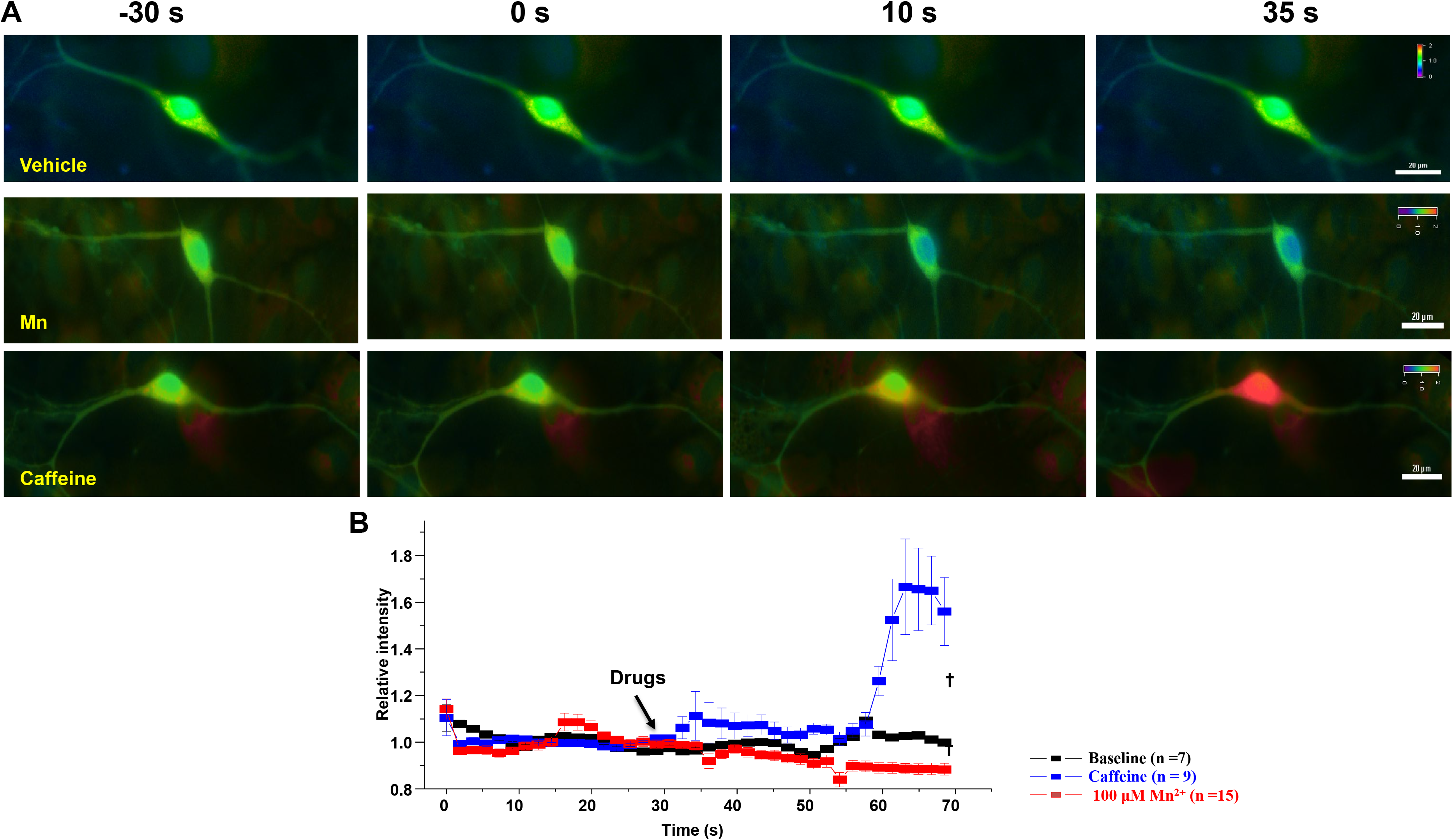
Mn does not increase intracellular calcium. Ratiometric analyses were performed on Fura-2-loaded dopamine neurons. ***A,*** Representative pseudo color images indicating the 340/380 ratio via a visible spectrum heat map (violet – minimum Ca^2+^; red – maximum Ca^2+^). Left panel shows representative 340/380 ratio images before and after drug or vehicle application, right panel shows identical view-fields 30-seconds after vehicle (top), 100 μM Mn (middle) and 20mM caffeine (bottom). ***B,*** Mean fluorescence intensities for each condition (normalized to the initial 30 s intensity). Data are plotted in 1.5 s interval for vehicle (black squares), Mn^2+^ (red squares), caffeine (blue squares). Scale bar: 20 μm. †: p < 0.01.

**Supplemental Figure 3.**
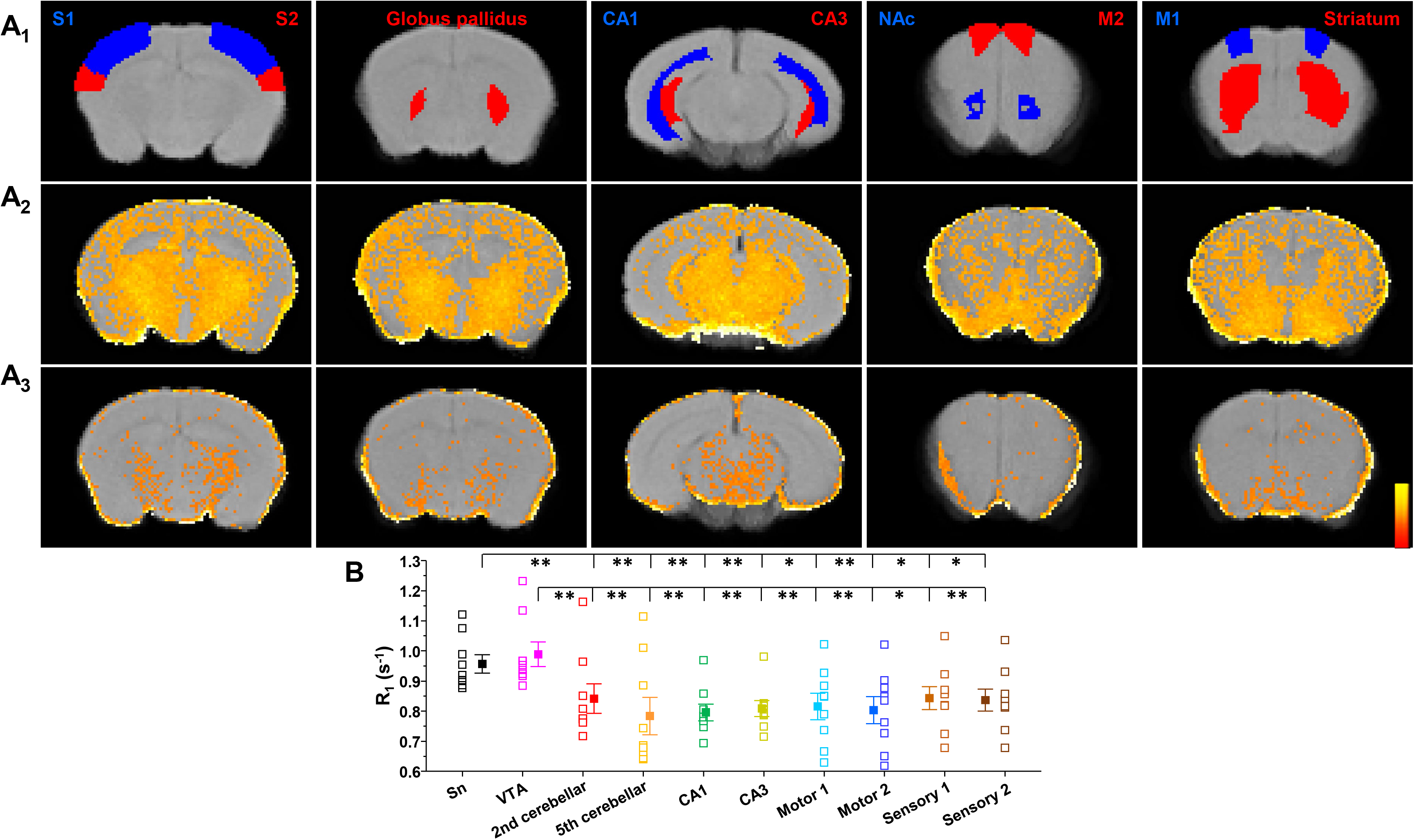
Ca^2+^ channel blockade using nifedipine reduces the effects of Mn^2+^ on T_1_ relaxation in several brain regions. ***A_1_*,** Mouse brain atlas-based segmentation of the primary (S1), secondary (S2) somatosensory cortex, globus pallidus, CA1, CA3, substantia nigra compacta (SNc), primary motor cortex (M1), secondary motor cortex (M2), and striatum. ROI, region of interest. ***A_2_***, Effects of Mn^2+^ accumulation on increasing brain T_1_ relaxation rate (R_1_ in msec^-1^). ***A_3_***, nifedipine treatment reduces the effects of Mn^2+^ accumulation by reducing its concentration in brain. Thus, maps of mice treated with nifepidine had a slower rate of T_1_ relaxation, or lower R_1_ in the case of the shown parametric maps (and lower intensity). Scale bar indicates intensity of R_1_. ***B,*** A greater Mn^2+^ accumulation produces faster rates of T1 relaxation (R_1_) in VTA and SN than in other segmented cortical and subcortical nuclei (*F*_(9,72)_ = 3.96, *p* < 0.001, one-way ANOVA followed by Tukey’s test, *n* = 8 – 9 / group). **p < 0.01.

Supplemental movie 1: Representative live cell imaging of FFN200 release before and after MnCl_2_.

Supplemental movie 2: Representative live cell imaging of Ca^2+^ mobilization before and after MnCl_2_.

Supplemental movie 3: Representative live cell imaging of Ca^2+^ mobilization before and after NMDA (positive control groups).

Supplemental movie 4: Representative live cell TIRF imaging before and after MnCl_2_.

